# Methyl-CpG Binding Protein 2 Reads Histone Methylation via an Aromatic Cage to Regulate Gene Expression and Chromatin Association

**DOI:** 10.1101/2025.05.10.653294

**Authors:** Jyotirmayee Padhan, Babu Sudhamalla

## Abstract

Methyl-CpG binding protein 2 (MeCP2) is a DNA methylation reader, which is highly expressed in the central nervous system. Loss of function mutation of MeCP2, which impairs DNA binding, causes Rett syndrome, while gain of MeCP2 function causes MeCP2 duplication syndrome. MeCP2 functions as both transcription activator and repressor but its full range of activity remains largely unknown. Recent study suggests that beyond DNA binding, MBD domain of MeCP2 also interacts with methylated lysine residues on histone H3 tail. In this study we aimed to characterize the methyllysine binding pocket of MeCP2-MBD. Structural analysis reveals the presence of an aromatic cage formed by five residues, namely, W104, F132, Y141, F142, and F155, where the trimethyllysine of H3K27me3 binds, as confirmed in docking studies. Alanine substitution of these residues abolished binding of MeCP2-MBD with its reported ligand H3K27me3 in ITC and pull-down experiments. Genomic analysis of publicly available ChIP-seq data reveals that MeCP2 localizes with canonical histone methylation marks like H3K4me3, H3K9me3, H3K27me3 and H4K20me3 and regulate critical biological and cancer related pathways. Promoters of MeCP2 regulated genes, show signature of H3K4me3 and H3K27me3. In the genomic level, mutation of aromatic cage residue W104A, significantly reduced chromatin occupancy of MeCP2 as demonstrated by ChIP-qPCR experiments. Furthermore, our qRT-PCR experiment, shows that this aromatic cage dependent binding is necessary for regulating expression of MeCP2 target genes. Overall, our study establishes MeCP2 as a histone methylation reader and highlights the functional significance of its histone binding capacity in chromatin association and gene regulation.

## Introduction

Methylation of DNA is a heritable epigenetic modification that involves the addition of a methyl group to the cytosine base in DNA (1). This modification is catalyzed by a group of enzymes known as DNA methyltransferases (DNMTs), which transfer a methyl group from S-adenosylmethionine (SAM) to the C5 position of the cytosine, resulting in the formation of 5-methylcytosine (5mC) (2). The primary site of DNA methylation in mammals is the CpG dinucleotide, which is found in short stretches throughout the genome. These CpG sites often form CpG islands, regions that are typically unmethylated (2). DNA methylation primarily has been associated with transcriptional repression. For example, transposable elements and viral sequences in the mammalian genome are heavily methylated to silence their expression, which could otherwise lead to genomic instability and mutations (3,4).

DNA methylation is recognized by specialized proteins, which are classified into three distinct families: MBD (methyl-CpG binding) proteins, UHRF (ubiquitin-like with PHD and RING finger) proteins, and zinc finger proteins (5). The MBD proteins were the first to be characterized, with MeCP2 being the founding member of this family. MeCP2 was initially identified in 1992 due to its ability to bind to methylated DNA (6), and was later found to be implicated in neurological disorder called Rett syndrome (7,8). MeCP2 is largely an intrinsically disordered protein, with only two domains exhibiting well-defined secondary structures: the MBD domain and the transcriptional repressor domain (TRD), which are connected by an intermediary region (9,10) (Figure 1A). The core structure of MBD domain folds into a wedge-like shape consisting of two short α-helices and three anti-parallel β-strands which are connected by a loop region (Figure 1B) (11). This domain recognizes the methylated CpG dinucleotide through two conserved symmetrical arginine residues (R111 and R133), forming a stair-shaped motif that interacts with DNA via hydrogen bonding, cation-π interactions, and nucleotide stacking (12,13). Conversely, the TRD of MeCP2 recruits histone deacetylases (HDACs) and Sin3A co-repressor complex to maintain a repressive state of the chromatin (14,15). Loss of DNA binding ability of MeCP2 manifests in Rett syndrome, where some of the critical DNA binding residues are mutated, like R133C, R168X, R255X, R270X, R294X, R106W, T158M, and R306C (16–20).

**Figure 1.**
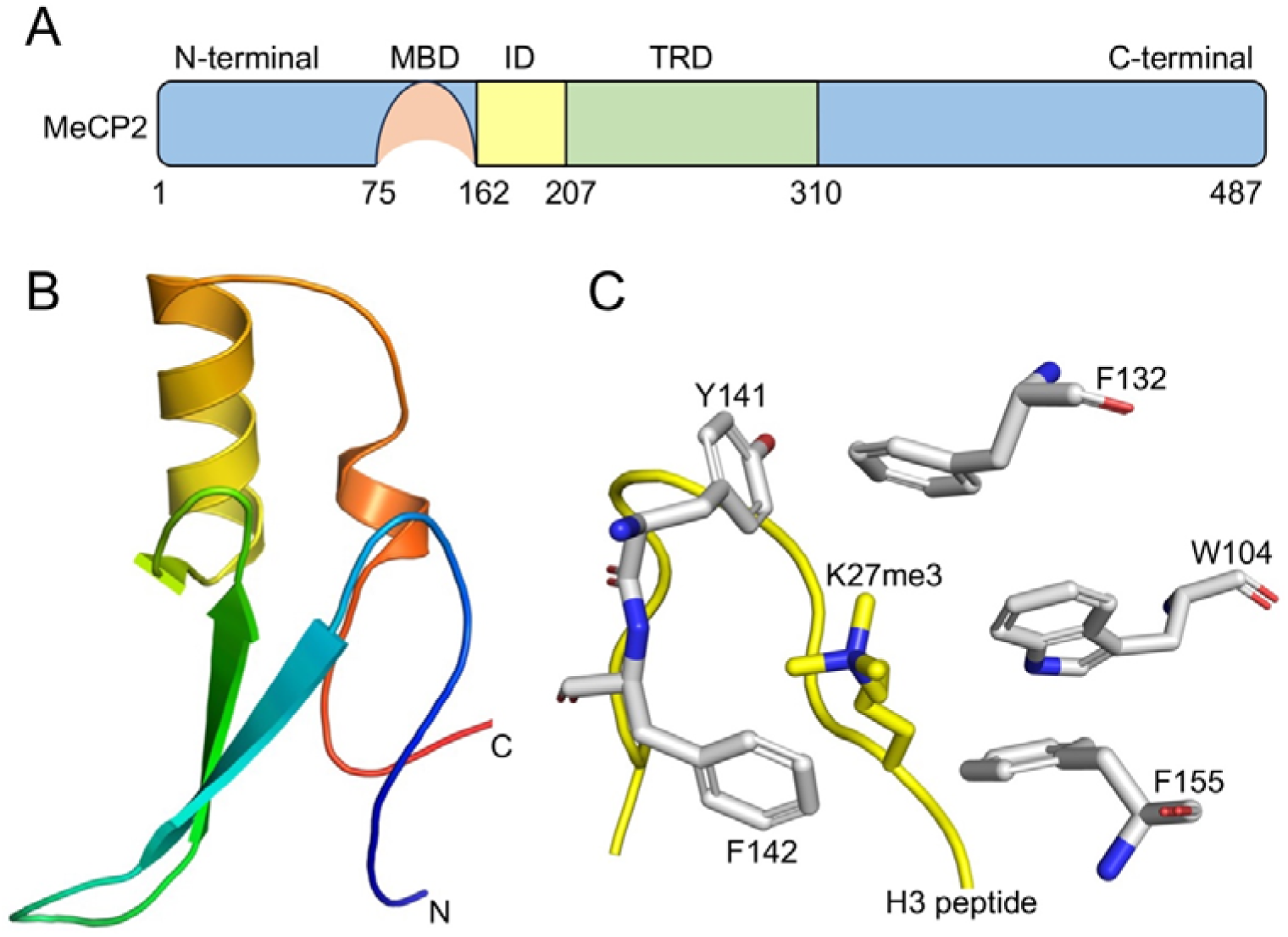
Domain organization of MeCP2 and docked structure of its MBD domain with H3K27me3 peptide. (A) Schematic representation of MeCP2 showing methyl-CpG binding domain (MBD) and a transcriptional repressor domain (TRD) connected by an intermediary domain (ID). (B) Cartoon representation of the X-ray crystal structure of MeCP2-MBD domain (PDB code 6OGK). (C) Close-up view of the docked complex of MeCP2-MBD domain and H3K27me3 peptide. The aromatic cage residues within the binding pocket are highlighted.

MeCP2 protein is abundantly expressed in nervous system and its transcription regulation is tightly controlled (21–23). Loss of MeCP2 function results in Rett syndrome, while duplication of its locus leads to its overexpression, causing MeCP2 duplication syndrome (24). Studies have indicated that MeCP2 functions not only as a transcriptional repressor but also plays a role in gene activation, depending on its interacting cofactors. In the mouse hypothalamus, for example, MeCP2 binds to the transcription activator CREB1 at the promoter of active genes (25). More recently, it was demonstrated that MeCP2 directly interacts with RNA polymerase II (RNA Pol II) at the transcription start site (TSS) of many active genes in human neural cells (26).

Additionally, various post-translational modifications (PTMs) also regulate MeCP2’s function by influencing DNA binding and its interactions with chromatin remodelers. For instance, acetylation of lysine 171 of MeCP2, reduce its affinity for HDAC1 and ATRX (27). Phosphorylation of MeCP2, triggered by calcium influx into neurons, facilitates the release of MeCP2 from the BDNF promoter (28,29). The functional versatility of MeCP2 contributed by PTMs regulate neuronal plasticity and morphology, cellular adaptation to stimuli, circuit formation, and the behavioral phenotype in mice (30).

Despite over three decades of MeCP2’s discovery, our understanding of this multifaceted protein’s role in gene regulation remain incomplete. Emerging evidence suggests that the MBD domain of MeCP2 not only recognizes methylated DNA but also interacts with specific histone methylation marks. Recent studies have shown that the MBD domain of MeCP2 binds to trimethyllysine marks on histone H3, such as H3K4me3, H3K9me3, H3K27me3, and H3K36me3 (31,32). However, the molecular mechanism by which a canonical DNA binding domain interacts with methylated histones remain unexplored. In this study, we focused on characterizing the methyllysine binding site within the MBD domain of MeCP2. Our findings reveal the presence of an aromatic cage within the MBD domain of MeCP2, where methyllysines are typically known to bind. We demonstrate that mutations of this aromatic cage residues disrupt the binding of MeCP2-MBD to its methylated histone ligand, H3K27me3, and significantly impair MeCP2’s localization to its target genes. These findings provide novel insights into the dual role of MeCP2 in chromatin recognition and regulation, thereby advancing our understanding of its function in gene expression.

## Results and discussion

### Recognition of histone H3K27me3 mark by MeCP2 via an aromatic cage in the MBD domain

Histone lysine methylation is a critical epigenetic mark that is dynamically regulated by lysine methyltransferases (KMTs) and lysine demethylases (KDMs) (33). The readers of histone lysine methylation are found in diverse protein families; however, the mechanism of methyllysine recognition remains largely conserved across these readers (34). Histone methylation readers typically feature an aromatic cage consisting of two to four aromatic residues, where the methyllysine binds (34–36). This binding is driven by cation-π interaction between the electron rich aromatic residues in the reader and the positively charged methylammonium group of lysine, along with hydrophobic and van der waals interactions (34–36). Recent studies have shown that the MBD domain of MeCP2, in addition to its well-known role in methyl-CpG binding, also interacts with the histone H3 trimethylated at lysine 27 (H3K27me3) mark (31). However, the precise binding site for this interaction remains unclear.

To identify the region of trimethyllysine binding within the MBD domain of MeCP2, we carefully examined its crystal structure (PDB code 6OGK) to search for the presence of such aromatic cage. We found that the MBD domain of MeCP2 contains five aromatic residues, viz, W104, F132, Y141, F142, and F155 that are arranged in a cage-like structure, which we hypothesized to be the possible site of methyllysine binding. To investigate the potential binding mode of MeCP2-MBD with H3K27me3 peptide, we performed protein-peptide docking using the HADDOCK2.4 web server. The five aforementioned aromatic residues were kept as active site residue and the H3(21–36)K27me3 was modeled in an extended conformation. HADDOCK generated 1000 docking models, which were clustered based on interface RMSD. The top-ranked cluster showed a favorable HADDOCK score of –53.76 ± 2.01, with the Kme3 group consistently occupying a hydrophobic pocket formed by aromatic cage residues (Figure 1C). The interaction was stabilized by cation–π interactions, hydrophobic, and salt bridge interaction between the trimethyllysine and residues in the binding pocket of the protein (Figure S1). These results suggest that the aromatic cage of MeCP2 MBD is a plausible Kme3-binding pocket.

### Mutation of aromatic cage residue disrupts binding of MeCP2-MBD with H3K27me3

To validate our docking results *in vitro*, we individually mutated these aromatic residues to alanine, overexpressed the MeCP2-MBD domain in *E. coli* cells, and purified the proteins to high homogeneity (Figure S2). We then performed ITC binding assays to assess the effect of these mutations on the binding affinity of MeCP2-MBD toward the H3K27me3 peptide. The wild-type MeCP2-MBD domain exhibited strong binding affinity for the H3K27me3 peptide, with a dissociation constant (*K*_D_) of 5.2 ± 0.4 µM (Figure 2A), which is consistent with previously reported value (32). In contrast, mutation of the aromatic cage residues, W104, F132, Y141, F142 and F155 to alanine drastically reduced the binding affinity for the H3K27me3 peptide, with a *K*_D_ ranging from >600 to >1000 µM (Figures 2B-F). Next, we assessed the binding affinity of MeCP2-MBD for the unmodified histone H3(15–36) peptide. The MeCP2-MBD domain bound weakly, with a dissociation constant (*K*_D_) of 37.5 ± 5.4 µM (Figure S3 and Figure S4-S5), indicating substantially reduced affinity compared with the H3K27me3(15–36) peptide.

**Figure 2.**
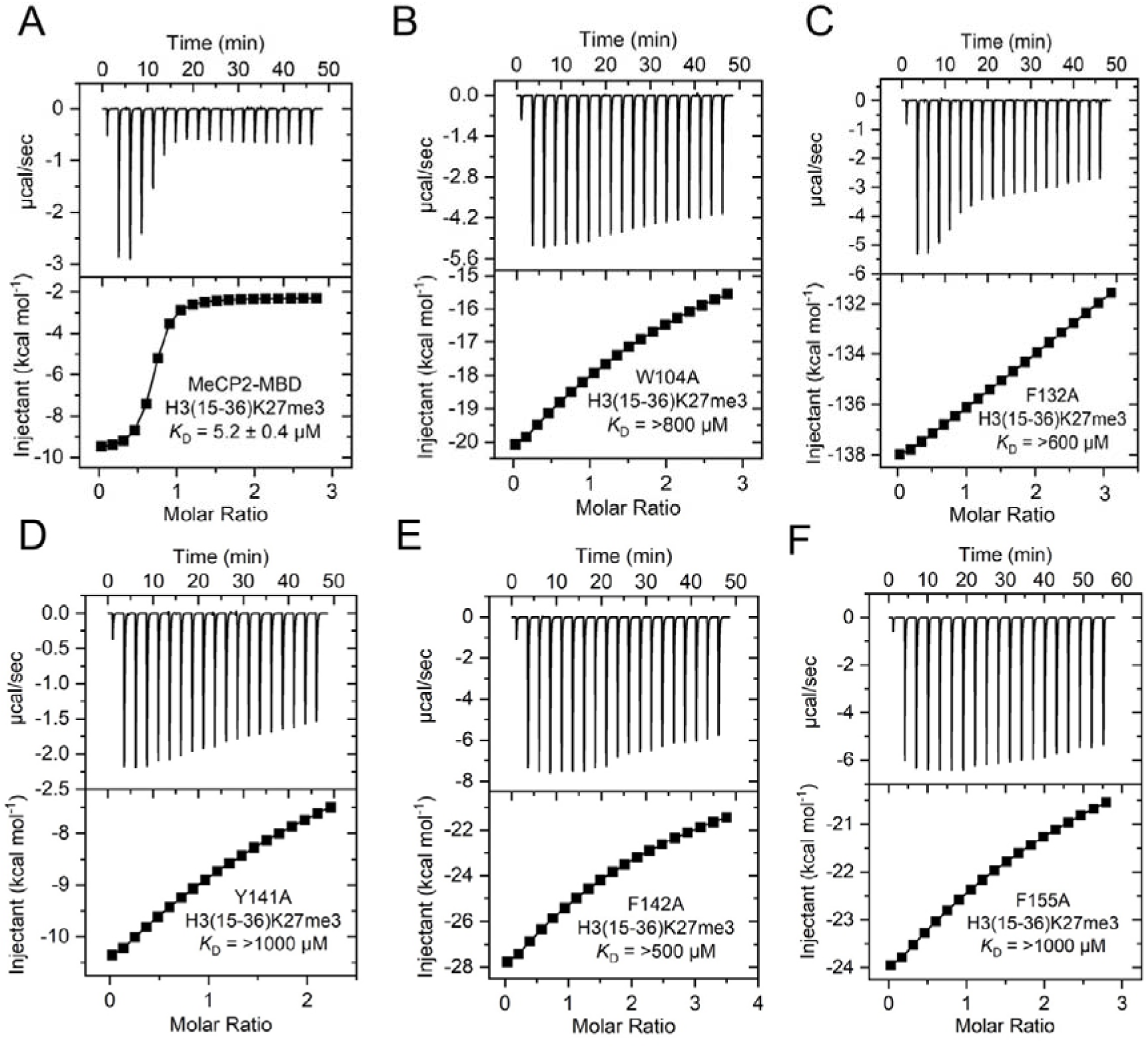
Exothermic ITC plots showing binding of MeCP2-MBD domain WT and mutants to H3(15–36)K27me3 peptide. (A) MeCP2-MBD WT, (B) MeCP2-MBD W104A, (C) MeCP2-MBD F132A, (D) MeCP2-MBD Y141A, (E) MeCP2-MBD F142A, and (F) MeCP2-MBD F155A. The calculated binding constants are indicated.

Next, we assessed the binding of MeCP2-MBD wild-type and its mutants to endogenous histones, which serve as more physiologically relevant substrates than peptides. Histones were isolated from HEK293T cells and incubated with 6xHis-tagged MeCP2-MBD wild-type and its mutant proteins (Figure 3A). Subsequent enrichment with Ni-NTA beads followed by western blotting with H3K27me3 antibody reveals that the wild-type MeCP2-MBD efficiently bound to H3K27me3 bearing histones, while mutants W104A, F132A, Y141A and F155A showed a complete loss of binding (Figure 3B). Notably, the F142A mutant exhibited a significant reduction in binding to H3K27me3 histones compared to wild-type protein (Figure 3B). Our findings from ITC binding and histone pull-down experiments align with previous reports on methyllysine readers. For example, the BAH domain of BAHD1, which also recognizes the H3K27me3 mark, features an aromatic cage consisting of tryptophan (W667) and two tyrosines (Y645 and Y669) (37). Mutations of these aromatic residues to alanine not only abolished the binding of BAH domain with H3K27me3 in ITC binding assays but also reduced the localization of BAHD1 with H3K27me3 in cells (37). Similarly, mutations of aromatic residues in other methyllysine readers, such as PHD, chromodomain, tudor domain, and MBT domain, have been shown to reduce ligand binding affinity (38–43).

**Figure 3.**
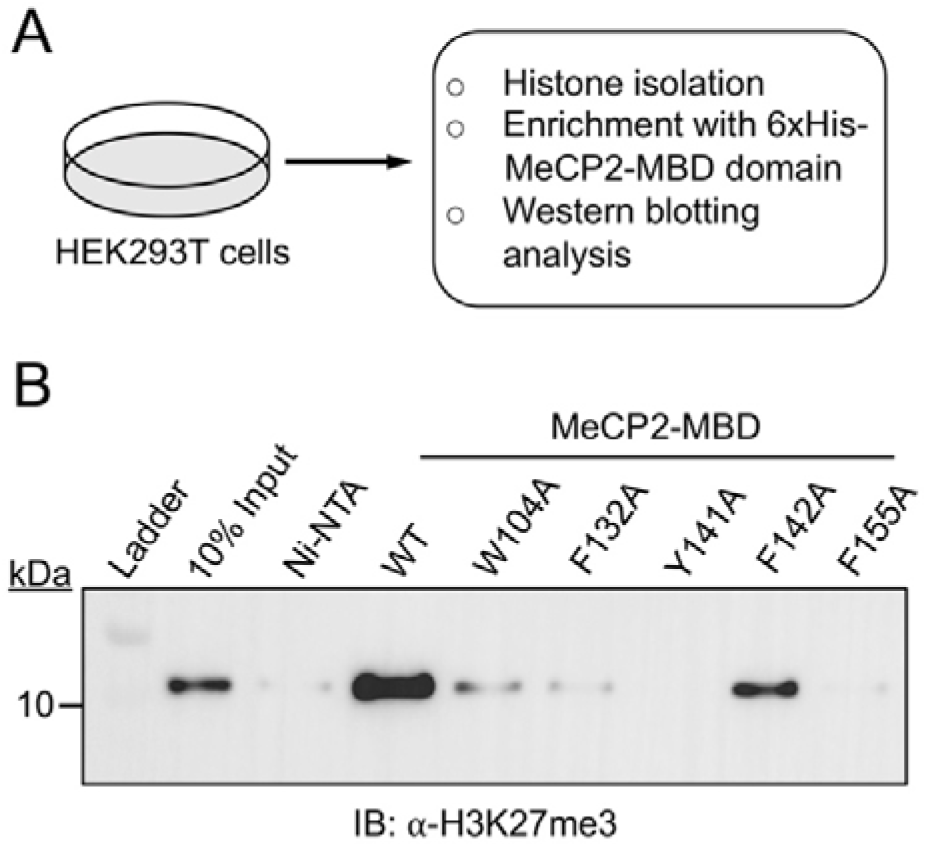
Interaction of MeCP2-MBD WT and mutants with endogenous histone H3. (A) Schematic depicting the preparation of endogenous histone from HEK293T cells followed by enrichment with MeCP2-MBD WT and mutant proteins. (B) Western blot analysis of histone H3K27me3 enrichment by MeCP2-MBD WT and mutant proteins.

Several reports suggest that some epigenetic readers also exhibit DNA binding capabilities, which promotes their engagement with chromatin and their affinity for DNA could be higher than for histones (44). For instance, the chromodomain of CBX8, a reader of the H3K27me3 mark and a component of polycomb repressive complex 1 (PRC1), binds to the H3K27me3 peptide with weak affinity (*K*_D_ = 0.7 mM) *in vitro*, while showing no interaction with its unmodified counterpart (45). However, electrophoretic mobility shift assays (EMSA) reveal that the CBX8 chromodomain binds both H3K27me3-modified and unmodified nucleosomes at similar concentration, suggesting that interactions with DNA primarily drive this binding (45). Given that, the MBD domain is also primarily a DNA interacting module, we propose that for MeCP2, the interaction of its MBD domain with DNA might be the main driving force behind its methyllysine recognition. Upon DNA binding, an induced-fit mechanism could occur, enabling the aromatic cage to open and accommodate the trimethyllysine. Our presumption aligns with observations that the binding pockets of methyllysine readers can undergo flexible structural changes to bind their cognate ligands (46,47).

### MeCP2-MBD interacts with DNA and histones through distinct interfaces within the same domain

The MBD of MeCP2 is a well characterized domain known to interact with DNA (6). Therefore, we next asked whether MeCP2-MBD engages DNA and histones through overlapping or distinct molecular interfaces. To investigate this, we first compared the structural models of MeCP2-MBD bound to DNA and to histone H3K27me3 peptide. Superimposition of the MeCP2-MBD-DNA crystal structure with the MeCP2-MBD-H3K27me3 docking model revealed that the two ligands occupy non-overlapping regions within the same domain (Figure 4A). While the DNA interacts with residues present on the surface of MBD, the H3K27me3 peptide binds to an adjacent but distinct interface containing the aromatic cage (Figure 4A). These structural observations suggests that MeCP2 can potentially accommodate both ligands simultaneously.

**Figure 4.**
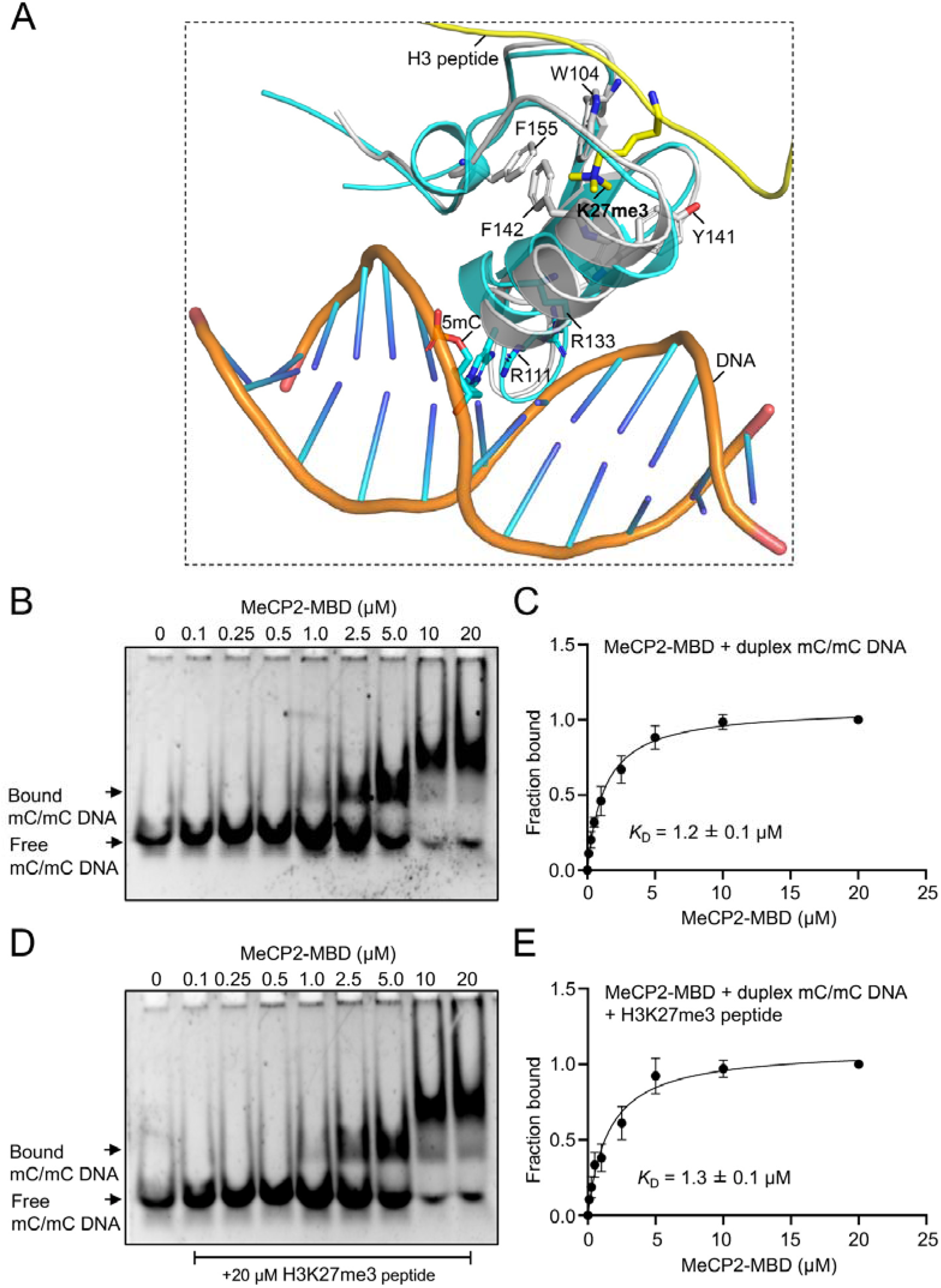
Dual recognition of methylated DNA and methylated histone H3K27me3 modifications by MeCP2-MBD domain. (A) Structural superposition of MeCP2-MBD-DNA complex with the docked model of MeCP2-MBD bound to H3K27me3 peptide, illustrating distinct binding interfaces for DNA and histone interactions. (B-E) EMSA titrations of the MeCP2-MBD with duplex mC/mC DNA in the absence and presence of H3K27me3 peptide. ImageJ was used to compute the fraction bound by measuring the raw intensity density measurements of each lane. Quantification of the gel images were done at least three independent experiments.

To experimentally validate this observation, we performed electrophoretic mobility shift assays (EMSA) using methylated *BDNF* DNA in the presence and absence of H3K27me3 peptide. The methylated *BDNF* DNA was incubated with increasing concentration of MeCP2-MBD and the reaction mixture was resolved in native polyacrylamide gels. The MeCP2-MBD efficiently bound methylated *BDNF* DNA with a dissociation constant *K*_D_ = 1.2 ± 0.1 μM (Figure 4B,C and Figure S6A). A similar *K*_D_ of 1.3 ± 0.1 μM was obtained, when the experiment was performed in the presence of H3K27me3 peptide (Figure 4D,E and Figure S6B) suggesting H3K27me3 peptide do not interfere with DNA-protein complex formation. Collectively, these findings confirm that histone interaction does not compete with DNA binding, supporting the structural model that MeCP2-MBD recognizes histones and DNA through spatially distinct binding interfaces.

### MeCP2 interacts and localizes with canonical histone methylation marks in the genome

Histone lysine methylation plays a crucial role in diverse nuclear functions, including heterochromatin formation, chromatin phase separation, and transcriptional regulation (33,48). The N-terminal tail of histones contains five canonical lysines that are subjected to methylation: H3K4, H3K9, H3K27, H3K36 and H4K20 (49,50). The occurrence of different methylation marks within the genome is indicative of distinct biological functions. For instance, the presence of H3K4me3 mark at gene promoters is associated with active transcription (51,52), while H3K9me3 is a hallmark of constitutive heterochromatin, and H3K27me3 mark is associated with facultative heterochromatin (53–55). These histone methylation marks are recognized by epigenetic reader proteins, which facilitate the recruitment of transcriptional regulatory complexes to the chromatin (36).

After characterizing the binding of MeCP2 with H3K27me3 mark, we next asked whether MeCP2 can also interact with other canonical histone methylation marks through its aromatic cage. To address this, we performed molecular docking of MeCP2-MBD domain with the trimethylated peptide corresponding to these canonical methylation sites. Docking study was performed in HADDOCK2.4 web server with parameters similar to those used for H3K27me3 docking. Our docking results demonstrates that the canonical histone methylation marks; H3(1-8)K4me3, H3(4-14)K9me3, H3(31-41)K36me3 and H4(14-26)K20me3 binds to the MeCP2-MBD with HADDOCK score of –59.6 ± 5.3, –70.9 ± 3.5, –69.4 ± 3.3 and –74.2 ± 3.6 respectively, with the trimethyllysine consistently positioned within the aromatic cage (Figure 5A-D). A recent study has also reported that MeCP2 binds to these histone methylation marks, including H3K4me3, H3K9me3, and H3K36me3, with dissociation constants (*K*_D_) of 0.12 µM, 0.075 µM, and 3.1 µM, respectively as determined in ITC experiments (32).

**Figure 5.**
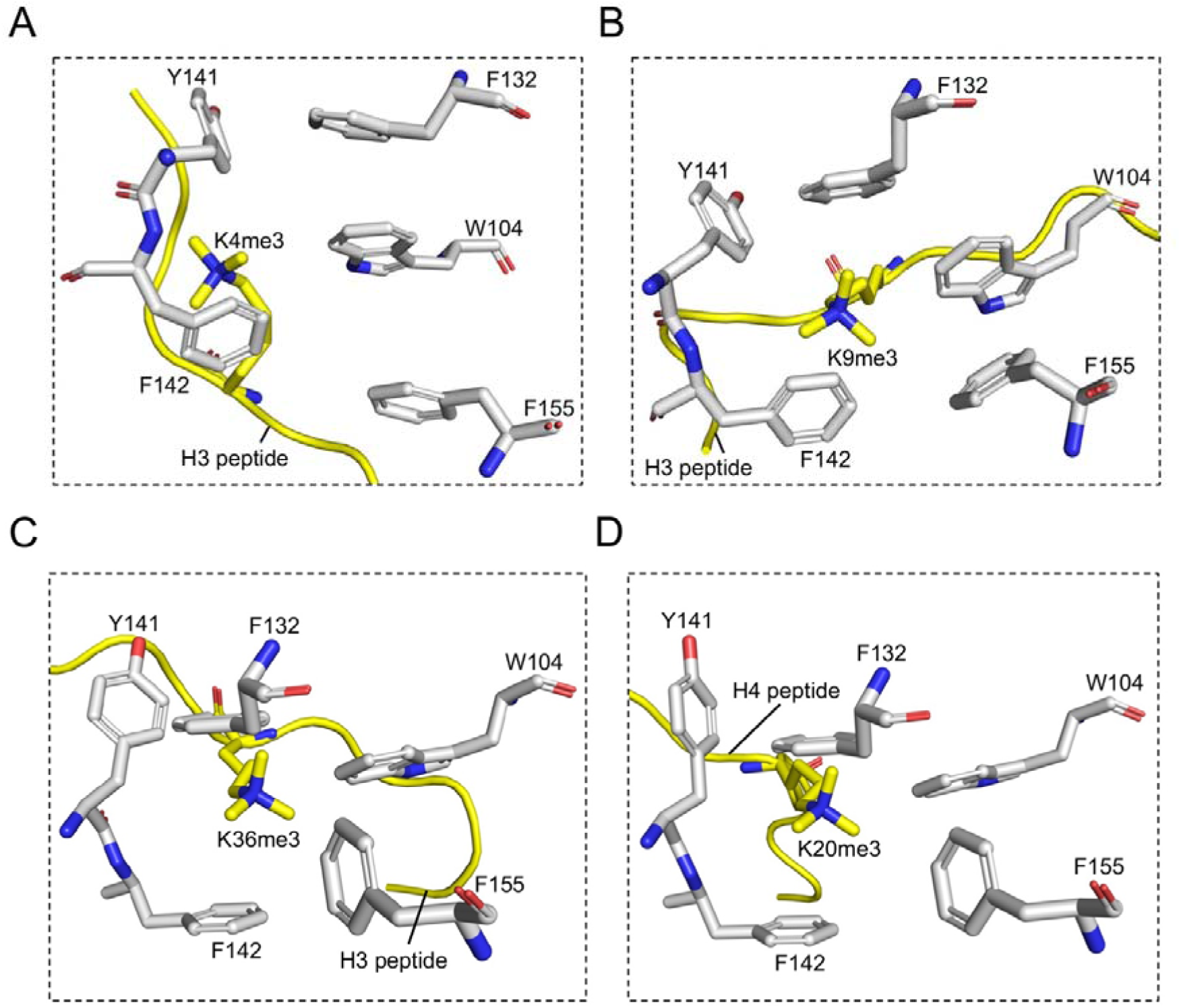
Close-up view of docked complex of MeCP2-MBD domain with (A) H3K4me3, (B) H3K9me3, (C) H3K36me3, (D) H4K20me3.

The strong interaction of MeCP2-MBD with canonical histone methylation marks, prompted us to investigate the colocalization of MeCP2 with these histone marks across the genome. We utilized publicly available ChIP-seq data to understand the localization of MeCP2 with the canonical histone methylation marks in the genome. Pearson correlation analysis showed that more than 50% of MeCP2 peaks overlap with peaks of H3K4me3, H3K9me3, H3K27me3, and H4K20me3 (Figure 6A). Among these marks, H3K4me3 exhibited the strongest positive correlation, with 66% of MeCP2 peaks coinciding with H3K4me3 peaks (Figure 6A). In contrast, despite MeCP2 binding to H3K36me3 (32), we observed a negative correlation between their genomic localizations (Figure 6A). Next, the significant peaks of MeCP2 and histone methylation marks were used to plot Venn diagram, in order to understand their overlap in the genome. Our Venn diagram analysis reveals that a total of 846 peaks (11.7%) out of 7216 peaks of MeCP2 overlaps with H3K4me3, H3K9me3, H3K27me3 and H4K20me3 peaks in the genome (Figure 6B). To further understand the colocalization of these canonical histone methylation marks in the MeCP2 occupied genes, we plotted heat maps. Heat map visualization demonstrate that transcription start site (TSS) of MeCP2 bound genes has abundant enrichment of H3K4me3 (Figure 6C). Additionally, the histone marks H3K9me3, H3K27me3, and H4K20me3 are distributed across the 5kb region from TSS of a significant subset of MeCP2 associated genes (Figure 6C). Consistent with the Pearson correlation analysis, the H3K36me3 marks are notably absent in MeCP2 occupied regions (Figure 6C).

**Figure 6.**
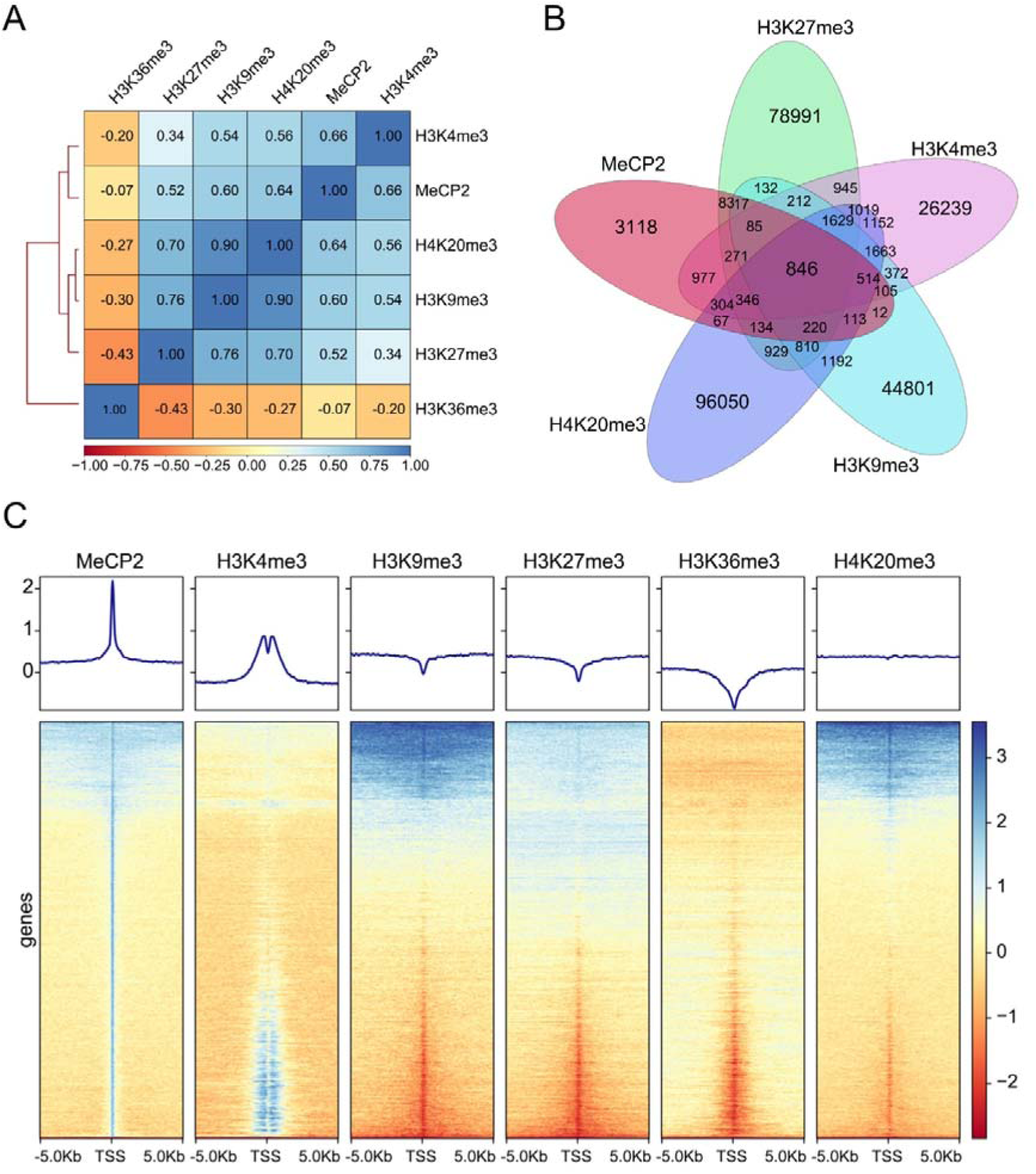
ChIP-seq analysis of MeCP2 localization and its association with canonical histone methylation marks in the genome. (A) Pearson correlation heat map depicting the relationship between MeCP2 binding and canonical histone methylation marks (B) Venn diagram illustrating the overlap of genome binding site between MeCP2 and histone marks H3K4me3, H3K9me3, H3K27me3 and H4K20me3. (C) Heat maps of ChIP-seq signals for MeCP2, H3K4me3, H3K9me3, H3K27me3, H3K36me3 and H4K20me3 within ±5 Kb of the TSS of MeCP2 bound genes (bottom). Line graphs showing the average ChIP-seq signal profiles around the TSS of MeCP2 bound genes (top). Genes were sorted from high to low based on MeCP2 peak intensity.

The localization of an epigenetic reader with specific histone modification signifies the readers role in regulating gene expression and chromatin dynamics (56). Initially identified as a DNA methylation reader, MeCP2 was thought to primarily act as transcription repressor by recruiting histone deacetylases (HDACs) and Sin3A repressor complex (14,15). However, subsequent studies have shown that, MeCP2 could also activate transcription by forming complex with the transcription factor CREB1 at the promoter of active genes (25). Our analysis supports this dual functionality of MeCP2, as it associates with both transcriptionally active marks, such as H3K4me3, and repressive marks, including H3K9me3, H3K27me3, and H4K20me3. The localization of MeCP2 with H3K9me3 and H4K20me3 can be attributed to its role in the formation and maintenance of pericentromeric heterochromatin (PHC) by deposition of H3K9me3 and H4K20me3, potentially by recruitment of specific histone methyltransferases (57). Consistent with this, earlier report also indicated that MeCP2 recruits histone methyltransferase specific for H3K9 methylation (58). And lastly, among all the canonical histone methylation marks, MeCP2 does not associate with H3K36me3 suggesting MeCP2 might not have any role in transcription elongation as H3K36me3 is a mark of transcription elongation and is present in the bodies of actively transcribed genes (59). Corroborating with our observation, it was noted that MeCP2 has no measurable effect in the rate of transcription elongation in mice brain (60).

### MeCP2 and histone methylation marks play crucial roles in regulating cellular pathways and cancer development

The binding of epigenetic readers to specific histone marks facilitates the recruitment of regulatory machinery, leading to the expression of relevant outputs that are implicated in various cellular functions. Since MeCP2 interacts with and localizes alongside several histone methylation marks, we sought to understand how this engagement influences biological pathways. To explore this, we conducted KEGG pathway enrichment analysis to assess the significance of histone marks recognized by MeCP2. The results, illustrated in the bubble map of KEGG enrichment analysis, demonstrate that MeCP2, in conjunction with canonical histone methylation marks, regulates critical cellular pathways such as Apelin, Wnt, Rap1, insulin, calcium, Hippo, cAMP, Ras, and mTOR signaling, as well as melanogenesis, autophagy, and cellular senescence (Figure 7). As MeCP2 is predominantly expressed in the nervous system, we also observed its involvement in neuronal pathways, including circadian rhythm regulation, glutamatergic and cholinergic synapse signaling, serotonergic synapse function, axon guidance, and neurodegenerative processes (Figure 7). Additionally, our analysis reveals that MeCP2 and histone methylation marks are implicated in the progression of various cancers, such as hepatocellular carcinoma, gastric cancer, breast cancer, colorectal cancer, chronic myeloid leukemia, and melanoma (Figure 7).

**Figure 7.**
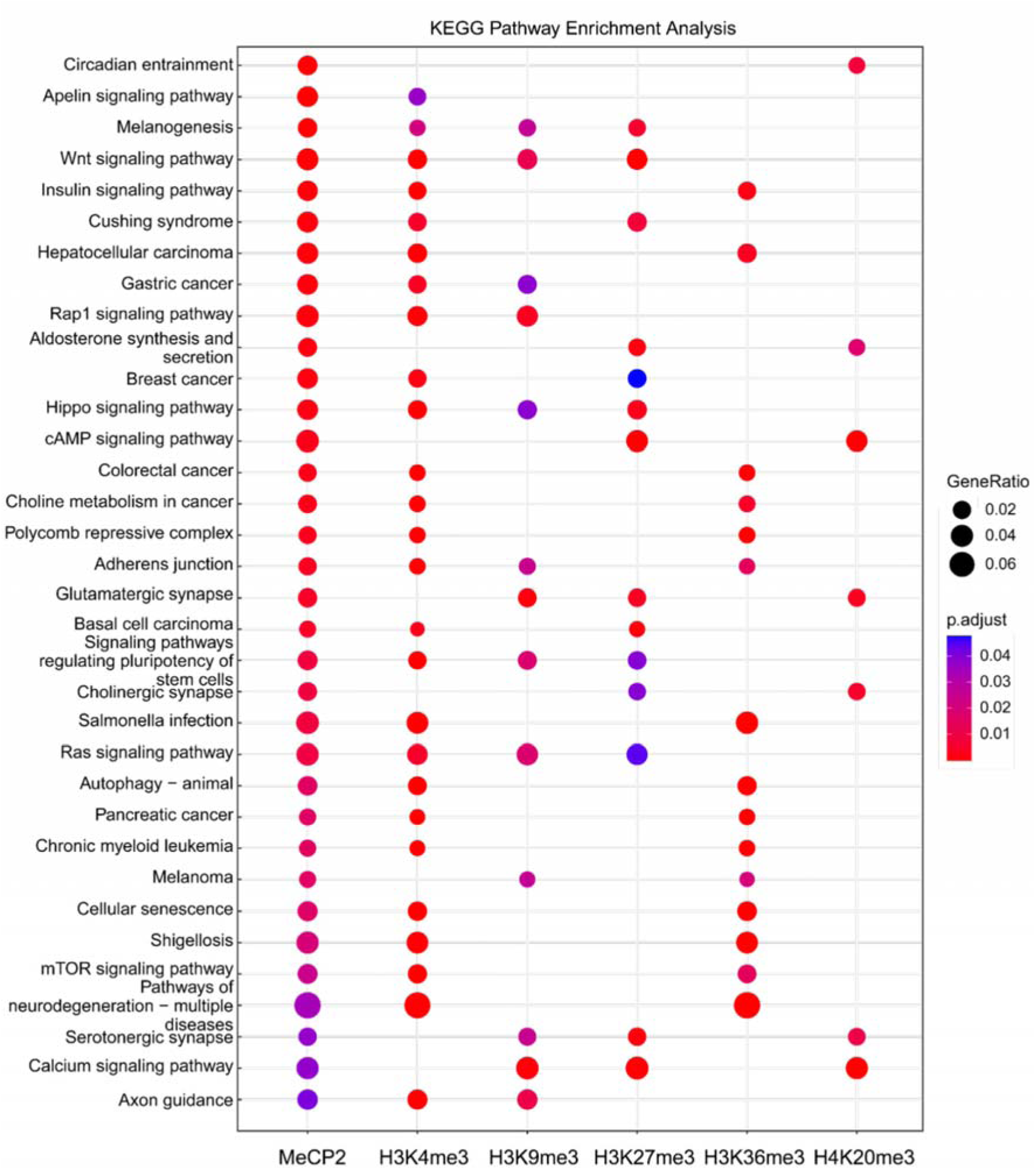
Bubble map of KEGG pathway enrichment analysis for MeCP2 and associated histone methylation marks. The Y-axis shows the names of the enriched pathways, while the size of each node indicates the gene ratio corresponding to MeCP2 and histone methylation marks H3K4me3, H3K9me3, H3K27me3, H3K36me3 and H4K20me3 marks. The *p*-value is represented by a color scale, with statistical significance increasing from blue (low significance) to red (high significance).

### Mutation in the aromatic cage of the MBD domain reduces MeCP2 localization to chromatin

Our ITC binding experiments and histone pull-down assays demonstrated that the MBD domain of MeCP2 recognizes trimethylated lysine 27 on histone H3 through an aromatic cage. Moreover, mutations in the aromatic cage of the MBD domain disrupt the binding of MeCP2 to both histone peptides and endogenous histones *in vitro*. Previously it has been reported that alanine mutations of tryptophan in the aromatic cage have abolished binding of methyllysine reader with its cognate ligand (61–63). Therefore, for rest of our experiments, we chose to focus on W104A mutant among the five aromatic cage residues of MBD domain of MeCP2. To investigate how MeCP2 interacts with chromatin in the genomic context, we mutated the aromatic cage residue W104 to alanine and transiently expressed HA-tagged wild-type MeCP2 and its mutant W104A in HEK293T cells (Figure S7). Given that MeCP2 is also a DNA methylation reader, we included the R133C mutation in our study. To validate the interaction between MeCP2 and histones at the cellular level, we performed immunoprecipitation assays. Our results show that HA-MeCP2 wild-type efficiently pulls down histone H3, while its mutant W104A shows a complete loss of binding to histone H3 in cells (Figure 8). In contrast, the R133C mutant retains its ability to bind histones (Figure 8), suggesting that residue R133 does not contribute to histone recognition.

**Figure 8.**
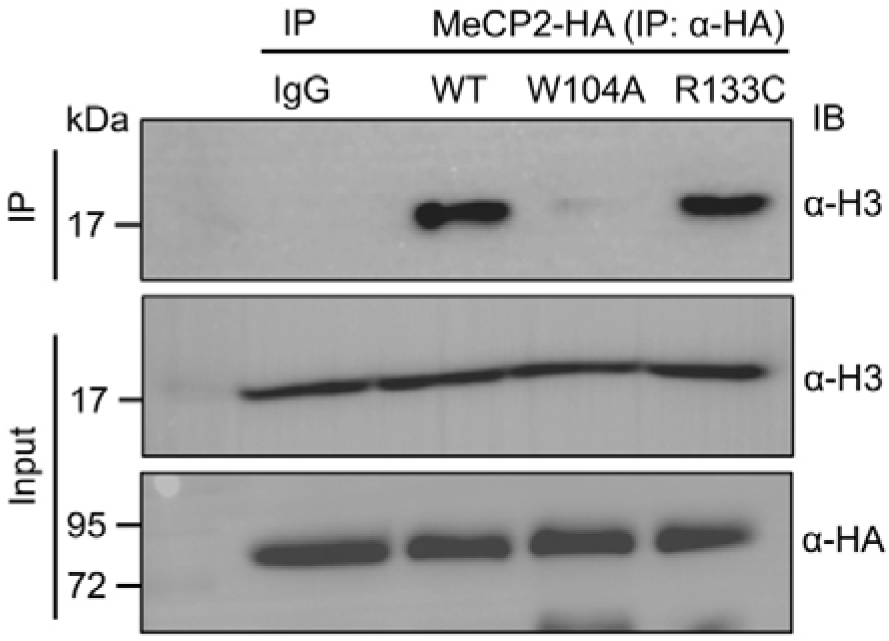
Immunoprecipitation (IP) of MeCP2 from HEK293T cells demonstrates that MeCP2 interacts with histone H3. Mutation of tryptophan at position 104 (W104A) disrupts the MeCP2– histone H3 interaction, whereas mutation of arginine at position 133 (R133A) does not impair histone H3 binding.

Next, we immunoprecipitated DNA from these cells and performed quantitative ChIP to assess MeCP2 localization at target genes. We selected ten genes previously reported as MeCP2 target; *DNMT1*, *SIRT1*, *ICAM1*, *HIPK3*, *DKK1*, *ILF3*, *ICAM3*, *KDM1A*, *ZNF713* and *HDAC1* (64). First, we examined the localization of MeCP2 and histone marks in the bodies of these genes. Analysis of publicly available ChIP-seq data reveals that MeCP2 and the histone marks H3K4me3 and H3K27me3 are enriched in the promoter of these aforementioned genes (Figure 9A and Figure S8). We validated this observation by performing ChIP-qPCR (Figure S9). Next, we performed ChIP-qPCR from immunoprecipitated DNA from cells expressing either WT or mutated MeCP2 plasmid. Our analysis of ChIP-qPCR experiment shows that mutation of aromatic cage residue W104 leads to more than 50% reduction in enrichment of MeCP2 in its target genes and this reduction is comparable to that observed for R133C mutation (Figure 9B). Together, these finding highlight the importance of aromatic cage of MeCP2 in histone recognition and chromatin localization. While R133C mutation is well established to impair DNA binding, the loss of histone interaction in case of W104A mutant suggest that histone binding also contributes to MeCP2’s genomic localization independent of DNA methylation. This dual mode of chromatin binding may be crucial for MeCP2’s function in the context of gene regulation.

**Figure 9.**
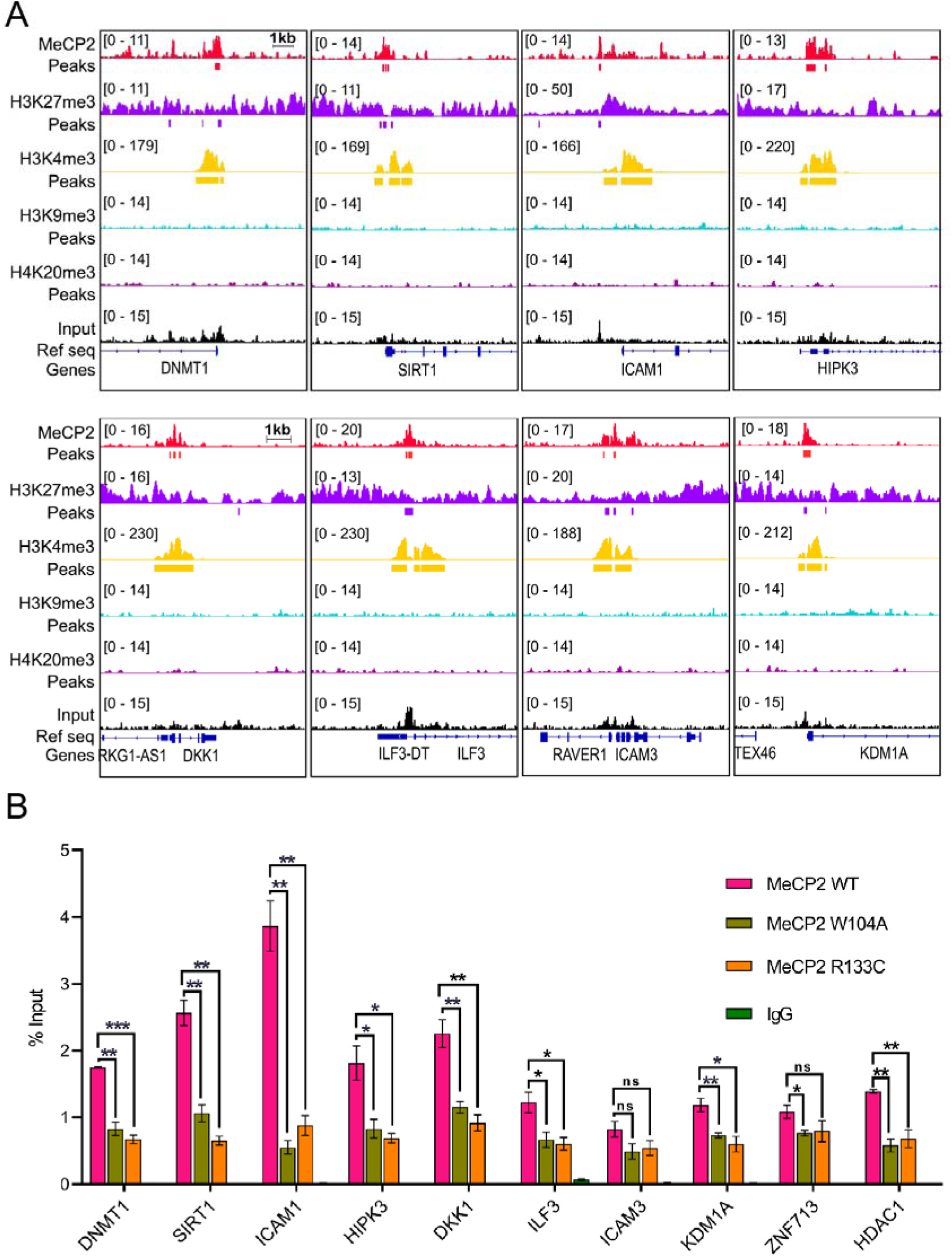
ChIP-seq analysis of MeCP2 binding to the human genome. (A) IGV snapshots showing the co-localization of MeCP2 and different histone methylation marks at the promoters of MeCP2 target genes. (B) ChIP-qPCR analysis showing reduced MeCP2 localization at target sites upon mutation of the aromatic cage residue (W104A) and the DNA-binding residue (R133C). Data are presented as mean ± SD (n = 3). Statistical significance was determined using Student’s t-test (\**p* ≤ 0.05; \*\**p* ≤ 0.01; \*\*\**p* ≤ 0.001; ns, not significant).

Our analysis also suggests that the promoter of some MeCP2 target genes contain both transcription activating H3K4me3 and transcription repressing H3K27me3 marks. The presence of both these histone modifications in a gene promoter is a signature of bivalent domain. Bivalent domains have traditionally been associated with expression of developmentally regulated genes in embryonic stem cells (ESC) keeping them in poised state (65,66). However, more recently it has been suggested that bivalency is a mechanism commonly used by cells to tightly regulate the expression of tissue specific genes (67). Hence, based on our observation, we propose that MeCP2 rather than functioning solely as a transcription activator or repressor, plays a broader transcriptional regulatory role.

### Mutation in the aromatic cage of the MBD domain alters expression of MeCP2 target genes

Our ChIP-qPCR analysis demonstrates that mutation of aromatic cage residue W104 significantly reduces chromatin occupancy of MeCP2, indicating that histone binding is critical for chromatin association. To access whether this binding is necessary for regulating the expression of MeCP2’s target genes, we performed qRT-PCR. We selected eight genes, *DNMT1, SIRT1, ICAM1, HIPK3, DKK1, ILF3, KDM1A,* and *HDAC1*, which showed significant reduction in MeCP2 localization upon the aromatic cage W104 mutation (Figure 9B). We transiently transfected HEK293T cells with HA-tagged MeCP2-WT, MeCP2-W104A and MeCP2-R133C plasmid and isolated RNA to perform qPCR. Our qPCR analysis shows that four MeCP2 target genes, *DNMT1, SIRT1, KDM1A,* and *HDAC1* exhibited significant reduction in expression in the presence of wild-type MeCP2 (Figure 10), which is consistent with MeCP2’s role as a transcription repressor. In contrast, W104A and R133C mutation does not reduce the expression of these genes similar to WT-MeCP2 (Figure 10). This finding suggests that, W104 and R133 residues which are critical for chromatin binding, are also required for MeCP2 mediated transcription repression. The remaining four genes does not show any significant changes in expression (Figure S10).

**Figure 10.**
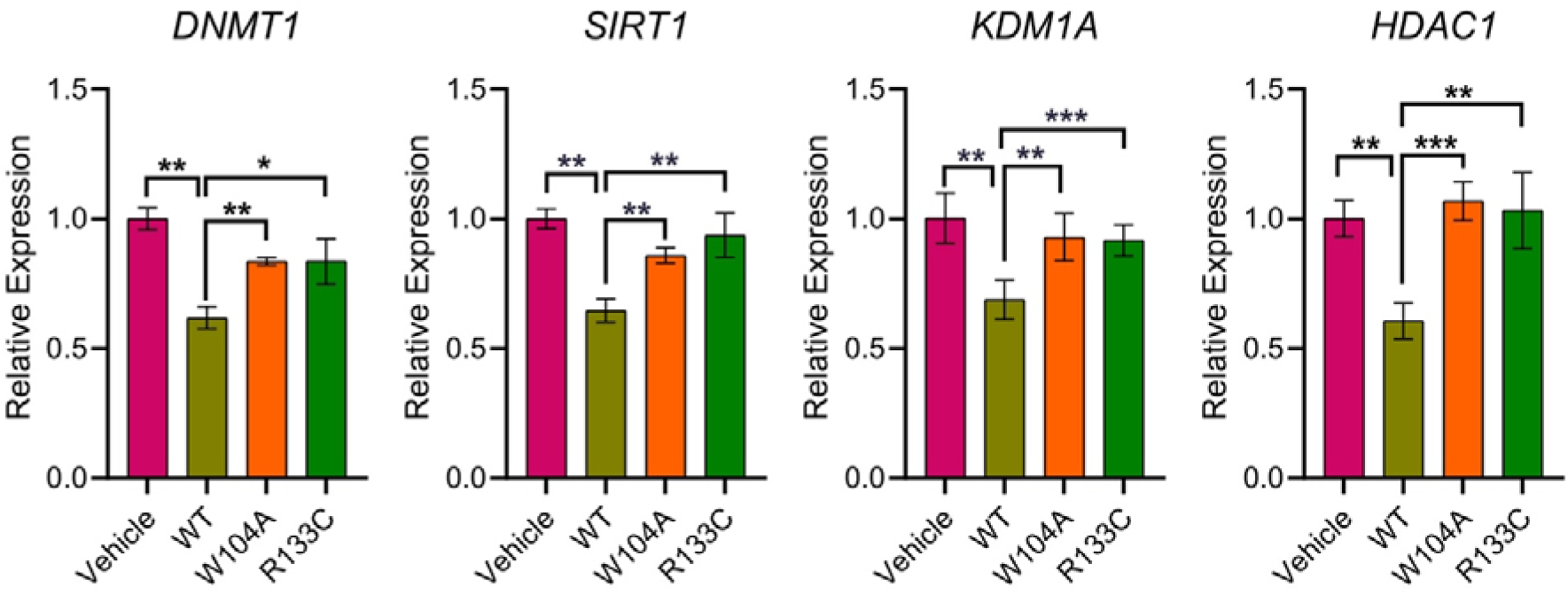
qRT-PCR analysis of MeCP2 target genes in the presence of MeCP2-WT, MeCP2-W104A and MeCP2-R133C. Data are presented as mean ± SD (n = 3). Statistical significance was determined using Student’s t-test (\**p* ≤ 0.05; \*\**p* ≤ 0.01; \*\*\**p* ≤ 0.001; ns, not significant).

## Conclusion

In conclusion, our study identifies MeCP2 as a histone methylation reader, highlighting its ability to bind methylated lysine residues in histones, specifically through an aromatic cage in its MBD domain. Mutation of key aromatic residues disrupts MeCP2’s binding to histone marks, such as H3K27me3, impairs its chromatin localization and alters expression of MeCP2 target genes. Genomic analysis further demonstrates that MeCP2 interacts with histone marks like H3K4me3 and H3K27me3 at gene promoters, regulating critical biological and cancer-related pathways. These findings underscore the importance of MeCP2’s histone-binding ability for its chromatin association and gene regulatory functions.

## Experimental procedures

### General materials, methods, and equipment

The plasmids for bacterial and mammalian expression were obtained from Addgene and Sino Biological, respectively. Mutagenesis primers, ChIP-qPCR primers and qPCR primers were obtained from Sigma-Aldrich (Table S1-S3). Commercially available competent bacterial cells were used for protein expression and mutagenesis. HEK293T cells were obtained from American Type Culture Collection and used following the manufacturer’s protocol. All of the antibodies used in this study were purchased from established vendors and used following the manufacturer’s protocol. The histone H3(15–36)K27me3 peptide was synthesized by S. Biochem Ltd and purified by HPLC to 98% purity (Figure S11). The concentrations of the peptides were determined on the basis of the observation that 1 mg/ml peptide generates an absorbance of 30 at 205 nm (68). The integrity of the purified peptides was confirmed by MALDI-MS (Figure S12).

### Protein-peptide docking

Protein–peptide docking was performed using the HADDOCK2.4 web server (69). The 3D structure of the MeCP2-MBD domain was obtained from the PDB (PDB code 6OGK), and the peptide representing H3(21-36)K27me3, H3(1-8)K4me3, H3(4-14)K9me3, H3(31-41)K36me3 and H4(14-26)K20me3 was built in an extended conformation using PyMOL. Both the protein and peptide structures were uploaded to the server, and the active site residues within the protein’s putative binding site were specified. The docking process was carried out in three stages: 1) rigid-body energy minimization (it0), 2) semi-flexible simulated annealing refinement (it1), and 3) final refinement in explicit solvent (water refinement). The resulting top clusters were ranked based on the HADDOCK score, a weighted sum of van der Waals interactions, electrostatics, desolvation energy, and restraint violations. The best-ranked cluster, characterized by the lowest average HADDOCK score, favorable interface RMSD, and a high buried surface area, was selected for further analysis. All models were visualized and assessed using PyMOL.

### Mutagenesis, expression and purification of the MeCP2-MBD domain

The C-terminal 6xHis-tagged MeCP2-MBD domain bacterial expression construct pNIC-CH was obtained from addgene (a gift from Cheryl Arrowsmith, addgene catalog no. 162259). MeCP2-MBD domain mutants W104A, F132A, Y141A, F142A, and F155A were generated using the QuikChange Lightning site-directed mutagenesis kit (Agilent catalog no. 210519), and the resulting mutations were confirmed by DNA sequencing. Protein expression and purification were performed using previously described methods with slight modifications (70,71). The wild-type MeCP2-MBD or its mutant plasmids were transformed into One Shot BL21 star (DE3) *E. coli* competent cells (Invitrogen, catalog no. C601003) using pNIC-CH kanamycin-resistant vector. A single colony was picked up and grown overnight at 37 °C in 10 mL of Luria-Bertani (LB) broth in the presence of 50 μg mL^−1^ kanamycin. The culture was diluted 100-fold and allowed to grow at 37 °C to an optical density (OD_600_) of 1.0. Protein expression was induced overnight at 17 °C with 0.5 mM isopropyl β-D-1-thiogalactopyranoside in an Innova 44 Incubator shaker (New Brunswick Scientific). Proteins were purified as follows. Harvested cells were resuspended in 15 mL of lysis buffer [50 mM HEPES (pH 7.5), 500 mM NaCl, 5 mM β-mercaptoethanol, 5% glycerol, 25mM imidazole, lysozyme, DNase, and 1:200 (v/v) Protease Inhibitor Cocktail III (Calbiochem)]. The cells were lysed by pulse sonication and centrifuged at 13000 rpm for 40 min at 4°C. According to the manufacturer’s instructions, the soluble extracts were subjected to Ni-NTA agarose resin (QIAGEN, catalog no. 30210). After 20 volumes of wash buffer [50 mM HEPES (pH 7.5), 500 mM NaCl, 5 mM β-mercaptoethanol, 5% glycerol, and 25 mM imidazole] had been passed through the column, proteins were eluted with a buffer containing 50 mM HEPES (pH 7.5), 500 mM NaCl, 5 mM β-mercaptoethanol, 5% glycerol, and 250 mM imidazole. Proteins were further purified by gel filtration chromatography (Superdex-75) using an AKTA pure FPLC system (GE Healthcare) in two different high and low salt concentration buffers; (low salt concentration, 50 mM HEPES (pH 7.5), 200 mM NaCl, and 5% glycerol) and (high salt concentration, 50 mM HEPES (pH 7.5), 500 mM NaCl, and 5% glycerol). Purified proteins were concentrated using an Amicon Ultra-10k centrifugal filter device (Merck Millipore Ltd.), and the concentration was determined using a Bradford assay kit (Bio-Rad Laboratories) with bovine serum albumin as a standard. The proteins were aliquoted and stored at −80 °C before use.

### Isothermal titration calorimetry (ITC) measurements

ITC binding experiments were performed as previously described (70,71). Briefly, experiments were carried out on a MicroCal PEAQ-ITC instrument (Malvern). Experiments were conducted at 15 °C while the samples were being stirred at 750 rpm. Buffers of protein and peptides were matched to 50 mM HEPES (pH 7.5), 200 mM NaCl, and 5% glycerol. Each titration comprised one initial injection of 0.4 μL lasting 0.8 s, followed by 18 injections of 2 μL lasting 4 s each at 2.5 min intervals. The initial injection was discarded during data analysis. The microsyringe (40 μL) was loaded with a peptide sample solution, and the peptide concentration varied from 1.5 to 2 mM. It was injected into the cell (200 μL), occupied by a protein concentration of 80-100 μM. All of the data were fitted to a single-binding site model using the MicroCal PEAQ-ITC analysis software to calculate the stoichiometry (*N*), binding constant (*K*_D_), enthalpy (*ΔH*), and entropy (*ΔS*) of the interaction. The final titration figures were prepared using OriginPro 2020 software (OriginLab).

### Mammalian cell culture and transfection

HEK293T cells were cultured in Dulbecco’s Modified Eagle’s Medium (DMEM) supplemented with 10% fetal bovine serum (FBS) and a penicillin/streptomycin cocktail, and maintained in a humidified incubator at 37°C with 5% CO2. Once the cells reached approximately 90% confluency, they were removed from the incubator, the medium was discarded, and the cells were washed with cold PBS. The collected cells were then rewashed with cold PBS, pelleted, and stored as a dry pellet at −80°C for subsequent histone extraction.

The mammalian expression construct pCMV3-C-HA-MeCP2-E1 (NCBI: NM_004992.4, 1-486 aa) was obtained from Sino Biological (Catalog no. HG12052-CY). The following point mutations W104A and R133C were introduced in pCMV3-C-MeCP2-E1 plasmid as described above. Cells were transfected at 80% confluency with a ratio of 1 μg DNA per 1 × 10^6^ cells for pCMV3-C-HA-MeCP2-E1, and its mutant plasmids in different cell culture dishes. A 2:1 ratio of Lipofectamine 2000 (Invitrogen, catalog no. 11668019) to DNA was used. First DNA and lipofectamine were incubated separately for 5 min at RT in Opti-MEM (Gibco, catalog no. 31985070). Then the DNA and lipofectamine were mixed together and incubated at RT for 20 min and then added to the cells. Six hours post transfection, media was replaced. Cells were collected after 24 to 36 h post transfection and frozen as a dry pellet at −80 °C before use.

### Acid extraction of histones

Histones were extracted from HEK293T cells using the acid-extraction protocol as previously described (72). Briefly, frozen cell pellets from 100 mm dish of HEK293T cells were resuspended in 1 ml of hypotonic lysis buffer (10 mM Tris-HCl pH 8.0, 1 mM KCl, 1.5 mM MgCl_2_, 1 mM DTT, 1 mM PMSF, and protease inhibitor cocktail) and incubated for 30 min on nutator at 4 °C. Samples were then centrifuged (10,000*g*, 10 min at 4 °C), and the supernatant was discarded entirely. The nuclei was resuspended in 800 μl of 0.4 N H_2_SO_4_, vortexed intermittently for 5 min, and further incubated at 4 °C on a nutator for overnight. The nuclear debris was pelleted by centrifugation (16000*g*, 10 min at 4 °C), and the supernatant containing histones were collected. The histones were precipitated by adding 264 μl trichloroacetic acid drop by drop to histone solution and the tube was inverted several times to mix the solutions and then the sample was incubated on ice for 30 min. Finally, histones were pelleted by centrifugation (16000*g*, 10 min at 4 °C), and the supernatant was discarded. The histones pellet was washed twice with ice-cold acetone, followed by centrifugation (16000*g*, 5 min at 4 °C), and the supernatant was carefully removed. The histone pellet was air-dried for 20 min at RT and subsequently dissolved in an appropriate volume of ddH2O and transferred into a fresh tube. The aliquoted histones were stored at −80 °C before use.

### Ni-NTA pull-down assays and western blotting

Pull-down experiments were performed by following the method previously described (70,71). First, 15-20 μg of acid-extracted histones isolated from HEK293T cells were mixed with 50 μM recombinant 6xHis-tagged-MeCP2-MBD wild-type and its mutant proteins (in high salt buffer) were incubated at room temperature on a nutator for 2 hours. Then, 50 μL of Ni-NTA agarose (QIAGEN, catalog no. 30210) was added to the reaction mixture and incubated for 45 min at room temperature on a nutator. The beads were then washed three times with 1 mL of wash buffer containing 50 mM HEPES (pH 7.5), 500 mM NaCl, and 25mM imidazole. Bound histones were eluted by incubating the beads in 20 μL of elution buffer [50 mM HEPES (pH 7.5) and 500 mM imidazole]. Equal volumes of eluted samples were separated on a 12% SDS-PAGE gel and transferred onto a 0.45 μm PVDF membrane at a constant voltage of 80 V for 1 h at 4 °C. The membrane was rinsed in TBST buffer [50 mM Tris (pH 7.4), 200 mM NaCl, and 0.1% Tween 20] and blocked for 1h at room temperature in 5% milk buffer prepared in TBST. Immunoblotting was performed overnight at 4 °C with the H3K27me3 (Cell Signaling, catalog no. 9733S) primary antibody. The membranes were washed with TBST buffer thrice at room temperature for 5 min each. The blots were then incubated with the HRP-conjugated secondary antibodies goat anti-rabbit IgG (Invitrogen, catalog no. 31466) with 5% non-fat dry milk, diluted 1:20000 in TBST. The membranes were rewashed with TBST buffer thrice at room temperature for 5 min each. Protein bands were visualized by chemiluminescence using Super Signal West Pico PLUS substrate (Invitrogen, catalog no. 34577) following the manufacturer’s protocol.

### Preparation and annealing of methylated DNA oligonucleotides

The DNA substrate used for EMSA was derived from the promoter region of brain-derived neurotrophic factor (*BDNF*), a known MeCP2 binding sequence (73). The methylated DNA sequence (top strand, 5′-GCCCTGGAA/iMe-dC/GGAACTCTTCTGGCC-3′; bottom strand, 5′-GGCCAGAAGAGTTC/iMe-dC/GTTCCAGGGC-3′), was synthesized by Integrated DNA Technologies (IDT). For annealing, equimolar concentration (100 μM each) of complementary strands were mixed in duplex buffer (100 mM potassium acetate, 30 mM HEPES, pH 7.5), heated to 94°C for 2 min and gradually cooled down to room temperature to form double-stranded DNA.

### Electrophoretic mobility shift assay

For EMSA titrations, 0.5 μM of *BDNF* duplex mC/mC DNA was incubated with increasing concentrations of MeCP2-MBD (0-20 μM) in the reaction buffer (50 mM HEPES pH 7.5, and 100 mM NaCl) with a total reaction volume of 20 μL. Parallel reactions were performed in the presence of 20 μM H3K27me3 peptide. Reaction mixtures were incubated for 1 h at 4°C, followed by addition of 2.5 µL loading dye to each sample. Samples were loaded onto a 6% native polyacrylamide gel and electrophoresis was performed in 1X TBE buffer at 80 V in 4°C. Gels were stained with SYBR Gold (Invitrogen, catalog no. S11494) for 15-20 min and imaged using the Syngene G:BOX Chemi-XRQ imaging system. ImageJ was used to compute the fraction bound by measuring the raw intensity density of each lane. Quantification of the gel images was done for at least three independent experiments. The binding constant (*K*_d_) values were determined using a non-linear regression equation:

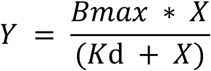

where *X* is the concentration of protein, *Y* is the fraction of DNA bound, and B*_max_* is the maximum binding in the same units as *Y* and *K*_d_ is the dissociation constant. The *K*_d_ values reported are averaged out of three experiments, and the error is calculated as the standard deviation between the three replicates.

### Public ChIP-seq data sets

MECP2 (SRR882078) (74), input DNA (SRR037637) and histone marks ChIP-seq raw data files of H3K4me3 (SRR639066) (75), H3K9me3 (SRR1617639) (76), H3K27me3 (SRR1617636) (76), H3K36me3 (SRR2877566) (77), and H4K20me3 (SRR1511591) (78), in IMR90 cells were obtained from Sequence Read Archive (SRA).

### Preprocessing and analysis of ChIP-seq data

ChIP-seq data were analyzed using the UseGalaxy server (79). Raw reads were quality-filtered and adapter-trimmed using the Trimmomatic trimming tool (80). Next, the trimmed reads were aligned in single-end mode with the human reference genome (hg19) using Bowtie2 (version 2.4.2) (81). Alignment followed by peak calling was performed with MACS2 (version 2.1.0) over input (82). As recommended for IDR (irreproducibility discovery rate) analysis, the final list of peaks was selected with an IDR threshold of 0.001 to obtain a conservative set of peaks. Peaks overlapping with blacklisted regions as defined by the ENCODE project were discarded. Heat maps and correlation plot were prepared using the deepTools2 (version 3.5.4) (83). Further analysis was performed in the R (version 3.4.0)/Bioconductor environment with plots generated relying on the ChIPseeker library (84). We used the Bioconductor package ChIPseeker (85) to display the overlaps between the different data sets and IGV for ChIP-seq signal visualization in a genomic context.

### ChIP-qPCR

ChIP-qPCR experiments were performed as previously reported with slight modifications (86). Briefly, the HEK293T cells were transfected with HA-tagged MeCP2-E1 wild-type and its mutants (W104A and R133C) plasmids in different cell culture dishes. Thirty-six hours post-transfection, cells were collected and fixed in 1% formaldehyde for 20 min, and quenched with 125 mM glycine for 5 min at RT. The cells were thoroughly washed with PBS supplemented with protease inhibitors. The pellets were resuspended in a high salt lysis/sonication buffer (25 mM Tris–HCl pH 7.5, 0.5% sodium deoxycholate, 0.1% SDS, 1% Triton X-100, 5 mM EDTA, 800 mM NaCl and protease inhibitors) and sonicated using Qsonica Q700 (pulsed 5s on, 30s off, for 60 s processing time). Insoluble cell debris was pelleted *via* centrifugation at 13,600*g* for 30 min at 4 °C. Dynabeads Protein G were incubated with 2.5 μg anti-HA antibody (Invitrogen, catalog no. 26183), or control IgG antibody (CST, catalog no. 5415) for 30 min at RT with rotation in Ab binding and washing buffer. The supernatant containing the soluble chromatin was incubated with Ab-conjugated magnetic beads at 4 °C for overnight, with rotation. Next, the beads were washed with wash buffer A (50 mM HEPES pH 7.9, 140 mM NaCl, 1 mM EDTA, 1% Triton X-100, 0.1% SDS and 0.1% sodium deoxycholate), wash buffer B (50 mM HEPES pH 7.9, 500 mM NaCl, 1 mM EDTA, 1% Triton X-100, 0.1% SDS and 0.1% sodium deoxycholate), and wash buffer C (25 mM Tris–HCl pH 7.5, 1 mM EDTA, 250 mM LiCl, 0.5% NP-40 alternative and 0.5% sodium deoxycholate). Finally, the beads were washed with tris-EDTA buffer for two times and eluted with elution buffer (10 mM Tris, pH 7.5, 1 mM EDTA, and 1% SDS) and incubated at 65 °C for 30 min. The eluted samples were treated with RNase A (Invitrogen, catalog no. EN0531) to a final concentration of 20 μg/ml and incubated at 65 °C for 8 h to digest any contaminated RNA. Next, the samples were treated with Proteinase K (Invitrogen, catalog no. 25530049) to a final concentration of 200 μg/ml and incubated at 45 °C for 2 h to digest proteins. Finally, the DNA was purified using a QIAquick PCR Purification Kit (QIAGEN, catalog. no. 28104) and used for qPCR analysis (Table S2).

### RNA isolation, reverse transcription, and quantitative real-time PCR

The HA-tagged MeCP2-E1 WT, MeCP2 W104A and MeCP2 R133C plasmids were overexpressed in HEK293T cells, and total RNA was extracted using the RNeasy mini kit (QIAGEN, catalog no. 74104) according to the manufacturer’s protocol. The isolated RNA was quantified using spectrophotometer. One microgram of total RNA and oligo dT primer was used to synthesize the cDNA using Verso cDNA Synthesis Kit (ThermoFisher scientific, catalog no. AB1453A) following the manufacturer’s protocol. Quantitative real-time PCR was performed in triplicate using the PowerUp SYBR Green Master Mix (ThermoFisher scientific, A25742) on a CFX96 touch real-time PCR detection system (Bio-Rad). Specific primers for each gene are collected from PrimerBank (87) for qRT-PCR experiments (Table S3). All mRNA quantification data were normalized to GAPDH, which was used as an internal control.

## Data availability

All data are included in the manuscript and supporting information.

## Acknowledgments

The authors thank infrastructural facilities supported by IISER Kolkata, and DST-FIST (SR/FST/LS-II/2017/93).

## Author Contributions

B.S. conceived the ideas. J.P., and B.S. designed the experiments. J.P. performed the experiments. J.P., and B.S. analyzed data. J.P., and B.S. prepared original draft. B. S. writing-reviewing and editing. B.S. supervision, B.S. resources. B.S. project administration. B.S. funding acquisition.

## Funding and additional information

This work was supported by grants from ANRF (CRG/2022/005242), SERB (EEQ/2020/000149), DBT Ramalingaswami Fellowship (BT/RLF/Re-entry/56/2018), and an intramural grant from IISER Kolkata (Ministry of Education), Government of India, to B.S. The authors also acknowledge the Prime Minister’s Research Fellowship (PMRF) program for providing doctoral fellowship and research grant to Jyotirmayee Padhan (PMRF ID-0501974), Government of India.

## Conflict of interest

The authors declare that they have no conflicts of interest with the contents of this article.

## Abbreviations

MeCP2: Methyl-CpG binding protein 2
MBD: methyl-CpG-binding domain
SAM: S-adenosylmethionine; 5mC, 5-methylcytosine
UHRF: ubiquitin-like with PHD and RING finger
TRD: transcriptional repressor domain.

## Notes

### Competing Interest Statement

The authors have declared no competing interest.

### Summary of Updates

Updated the introduction, methods sections and addded new results: -Docking of MeCP2-MBD with diferent methylated peptide like H3K4me3, H3K9me3, H3K36me3, and H4K20me3). -EMSA results

