## Supporting Information for "Methyl-CpG Binding Protein 2 Reads Histone Methylation via an Aromatic Cage to Regulate Gene Expression and Chromatin Association"

**Table of Contents**

1. Supplementary Tables S2
2. Supplementary Figures S3

**Supplementary Table S1.** List of the forward primers designed for site-directed mutagenesis. Reverse primers used are the reverse-complement to the given forward primers.

| **MeCP2-MBD Mutants** | **Primer Sequence** |
|---|---|
| W104A | CCCACCCTGCCTGAAGGCGCGACACGGAAGCTTAAG |
| F132A | CCCCAGGGAAAAGCCGCGCGCTCTAAAGTGGAGTTGATTG |
| Y141A | GTGGAGTTGATTGCGGCGTTCGAAAAGGTAGGCGACAC |
| F142A | GTGGAGTTGATTGCGTACGCGGAAAAGGTAGGCGACAC |
| F155A | CTGGACCCTAATGATGCGGACTTCACGGTAACTGGG |
| R133C | CCCCAGGGAAAAGCCTTTTGCTCTAAAGTGGAGTTGATTGCG |

**Supplementary Table S2.** List of primers used for the ChIP-qPCR.

| **ChIP-qPCR Primers** | | |
|---|---|---|
| **Gene** | **Forward** | **Reverse** |
| *DNMT1* | CAAAAGGGGAACCTTGTTCA | CCTGGGAGGAAGAAATAGGG |
| *SIRT1* | TAGACGCAACAGCCTCCG | GGCTGCGGGAGATTTAAACC |
| *ICAM1* | ATTGTCCGGGAAACTGGACG | ACAACAGGCGGTGAGGATTG |
| *HIPK3* | CCACTACCCCTCGCCCTA | CTGAGAGGAAACGGCGAAAC |
| *DKK1* | CGGTTCTCAATTCCAACGCT | CCCCTCTCACCTGGTAGTTG |
| *ILF3* | AGGCAGCTACTCCTACTCGAA | CACCACTTGTCCTCCTCCTAA |
| *ICAM3* | CATGGTCCAGTGGGAAAGGT | ATAGGCTTGTACGCCATCCC |
| *KDM1A* | AAACCCGAAAGTCCCTGGAG | GCAGCAAAGAACGTGTAGCT |
| *ZNF713* | AAAACAGACCCGGGAAAGCT | GACAGCCTCCCCGCAATG |
| *HDAC1* | ACTACTACGACGGTGAGCAC | CCTCCTCTCCGAGCCTCT |

**Supplementary Table S3:** List of primers used for the qRT-PCR.

| **qRT-PCR Primers** | | |
|---|---|---|
| **Gene** | **Forward** | **Reverse** |
| *DNMT1* | CCTAGCCCCAGGATTACAAGG | ACTCATCCGATTTGGCTCTTTC |
| *SIRT1* | TAGCCTTGTCAGATAAGGAAGGA | ACAGCTTCACAGTCAACTTTGT |
| *ICAM1* | ATGCCCAGACATCTGTGTCC | GGGGTCTCTATGCCCAACAA |
| *HIPK3* | TCACAAGTCTTGGTCTACCCA | CACATAGGTCCGTGGATAGTTTC |
| *DKK1* | CCTTGAACTCGGTTCTCAATTCC | CAATGGTCTGGTACTTATTCCCG |
| *ILF3* | AGCATTCTTCCGTTTATCCAACA | GCTCGTCTATCCAGTCGGAC |
| *KDM1A* | TGACCGGATGACTTCTCAAGA | GTTGGAGAGTAGCCTCAAATGTC |
| *HDAC1* | CGCCCTCACAAAGCCAATG | CTGCTTGCTGTACTCCGACA |

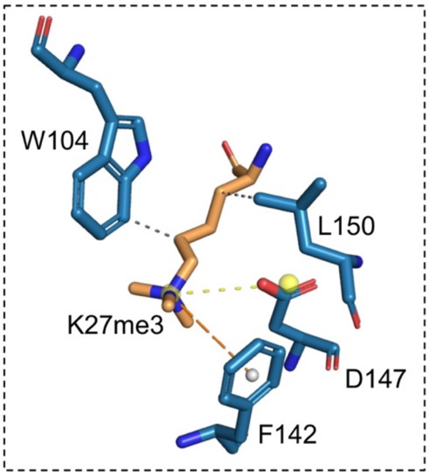


**Supplementary Figure S1.** Close-up view of H3K27me3 docked in MeCP2-MBD showing interactions between the protein and ligand: hydrophobic interactions with W104 and L150, cation-π interaction with F142 and salt bridge interaction with D147.

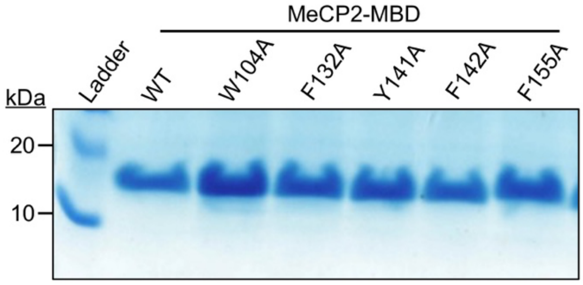


**Supplementary Figure S2.** Coomassie stained SDS-PAGE gel showing the overexpression of MeCP2-MBD wild-type and its mutants in BL21 Star (DE3) *E. coli* cells.

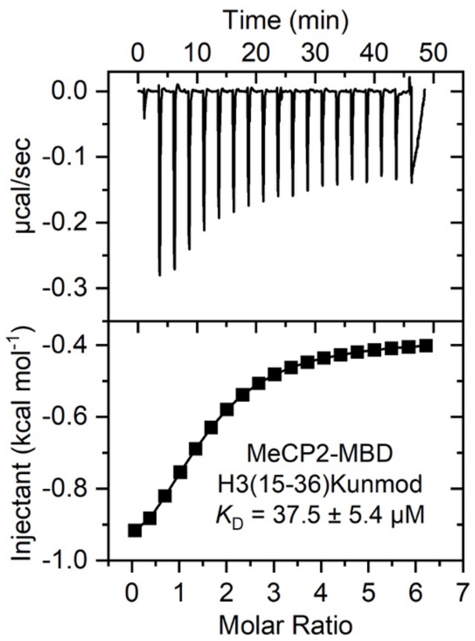


**Supplementary Figure S3.** Exothermic ITC plots showing binding of MeCP2-MBD domain to H3(15-36) unmodified peptide. The calculated binding constants are indicated.

### HPLC REPORT

| | | | |
|---|---|---|---|
| Sample: | Pep-685 APRKQLATKAARKSAPATGGVK | | Analyzed date: 21-03-2023 |
| Analyst: | Dr.AR-SBio | | |
| Column: | 4.6×250mm,Sinochrom ODS-BP 5μm | | |
| Solvent A | A: 0.1% Trifluoroacetic Acid in 100% Acetonitrile | | |
| Solvent B | B: 0.1% Trifluoroacetic Acid in 100% Water | | |
| Gradient: | | A | B |
| | 0.0min | 7% | 93% |
| | 25.0min | 32% | 68% |
| | 25.1min | 100% | 0% |
| | 30.0min | | Stop |
| Volume: | 5μl | | |
| Wavelength: | 220nm | | |
| Flow rate: | 1.0ml/min | | |

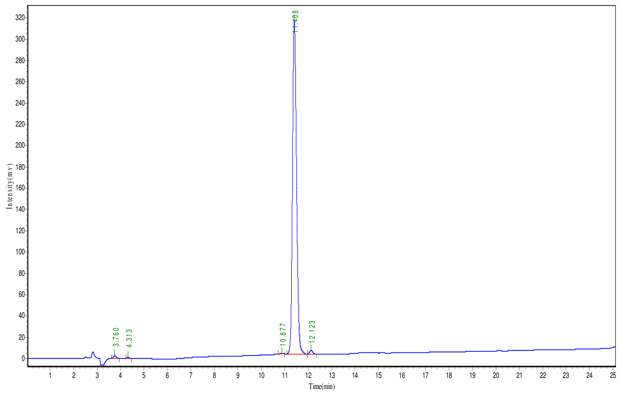


| Peak | Time | Height | Area | Conc. |
|---|---|---|---|---|
| 1 | 3.760 | 2243.948 | 19277.100 | 0.5076 |
| 2 | 4.313 | 1043.928 | 6693.750 | 0.1763 |
| 3 | 10.877 | 830.575 | 7119.000 | 0.1875 |
| 4 | 11.408 | 313463.625 | 3725986.500 | 98.1142 |
| 5 | 12.123 | 3722.384 | 38524.609 | 1.0144 |
| Total | | | | 100.000 |

**Supplementary Figure S4.** HPLC purity trace for the histone H3(15-36) peptide.

### MASS SPECTROMETRY REPORT

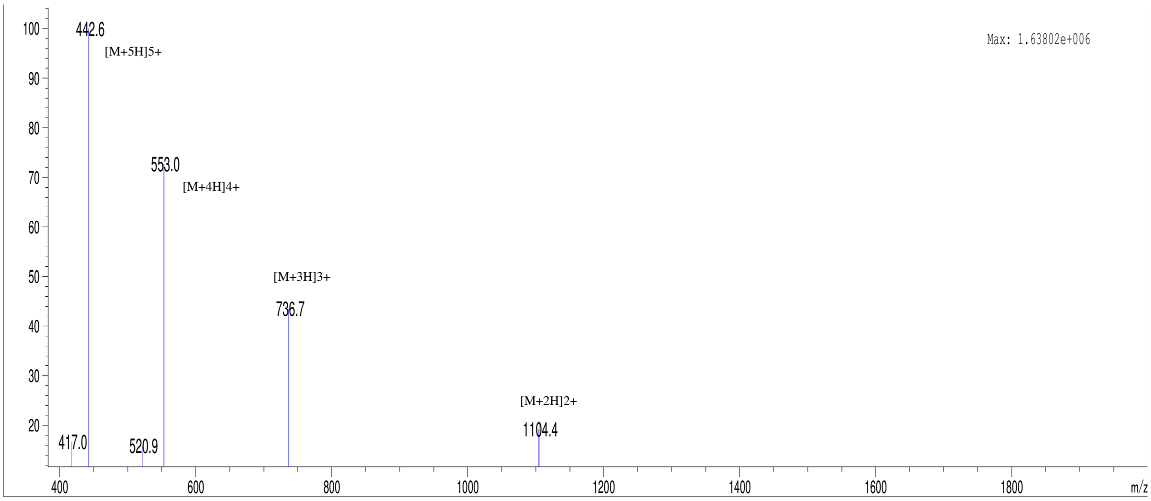


| Sample Description | | Instrument | Agilent-6125B | | |
|---|---|---|---|---|---|
| Analyzed date: | 22-03-2023 | Probe: | ESI | Probe Bias: | +4.5kv |
| Analyst: | Dr.AR-SBio | Nebulizer Gas Flow: | 1.5L/min | Detector: | 1.5kv |
| Sample: | Pep-685 APRKQLATKAARKSAPATGGVK | CDL: | -20.0v | T. Flow: | 0.2ml/min |
| M.W.: | 2207.57 | CDL Temp.: | 250 °C | B. Conc.: | 50%H2O/50%ACN |

**Supplementary Figure S5.** MS spectra for the histone H3(15-36) peptide.

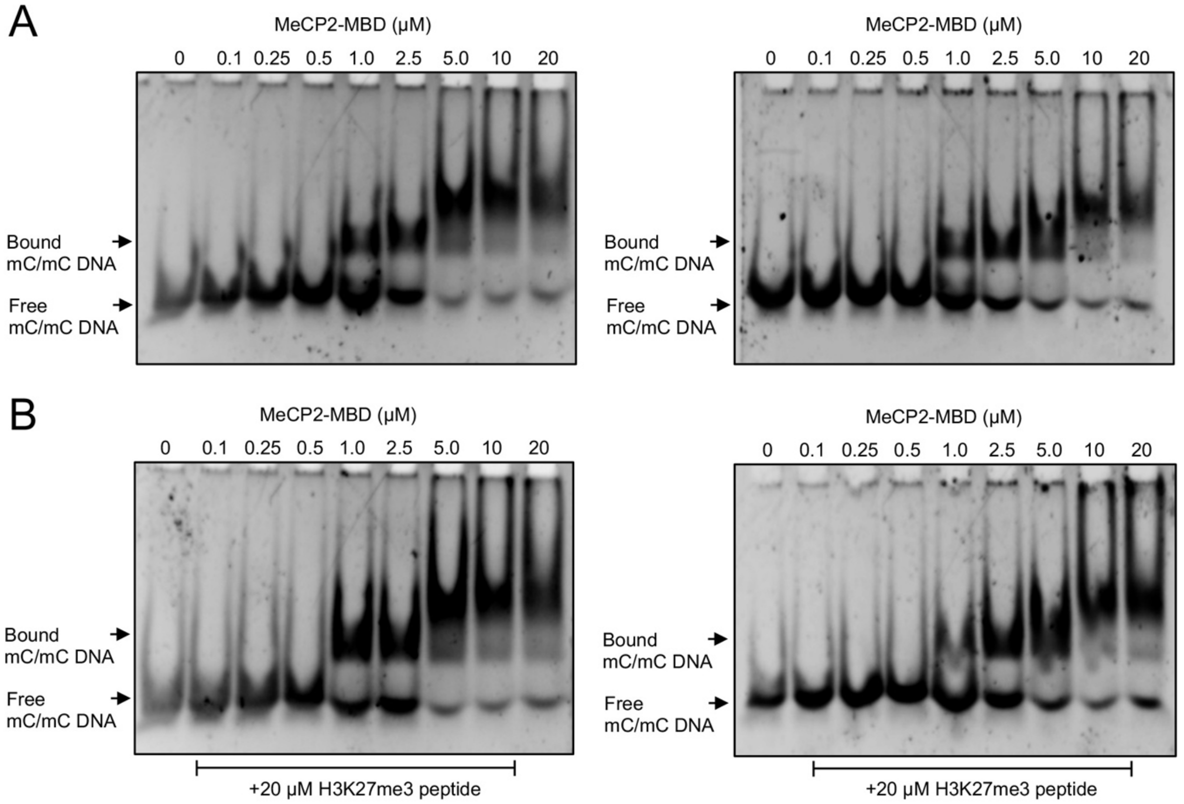


**Supplementary Figure S6.** EMSA titrations of the MeCP2-MBD with methylated DNA in the absence and presence of H3K27me3 peptide.

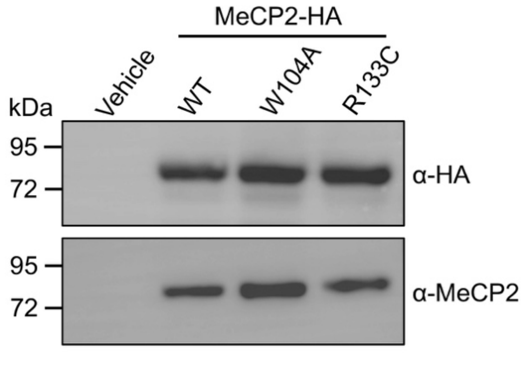


**Supplementary Figure S7.** Western blot analysis showing the expression of HA-MeCP2-WT and its mutants in HEK293T cells.

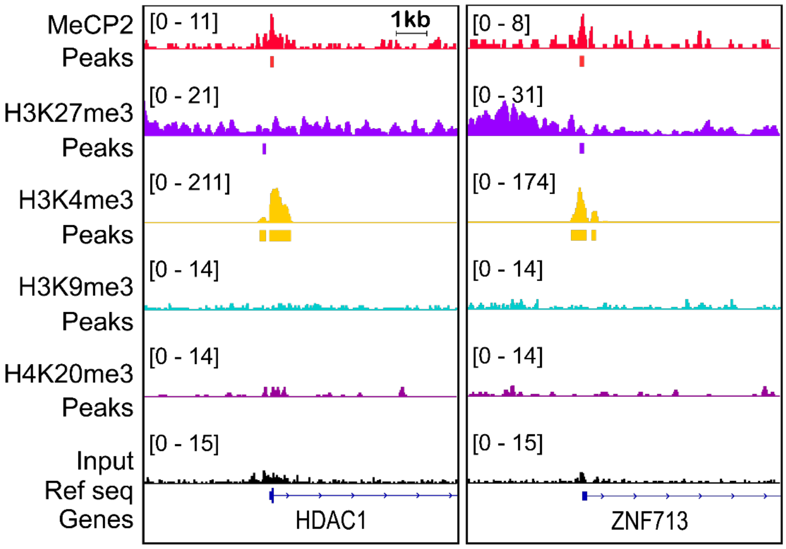


**Supplementary Figure S8.** IGV snapshots showing the co-localization of MeCP2 and different histone methylation marks at the promoters of MeCP2 target genes.

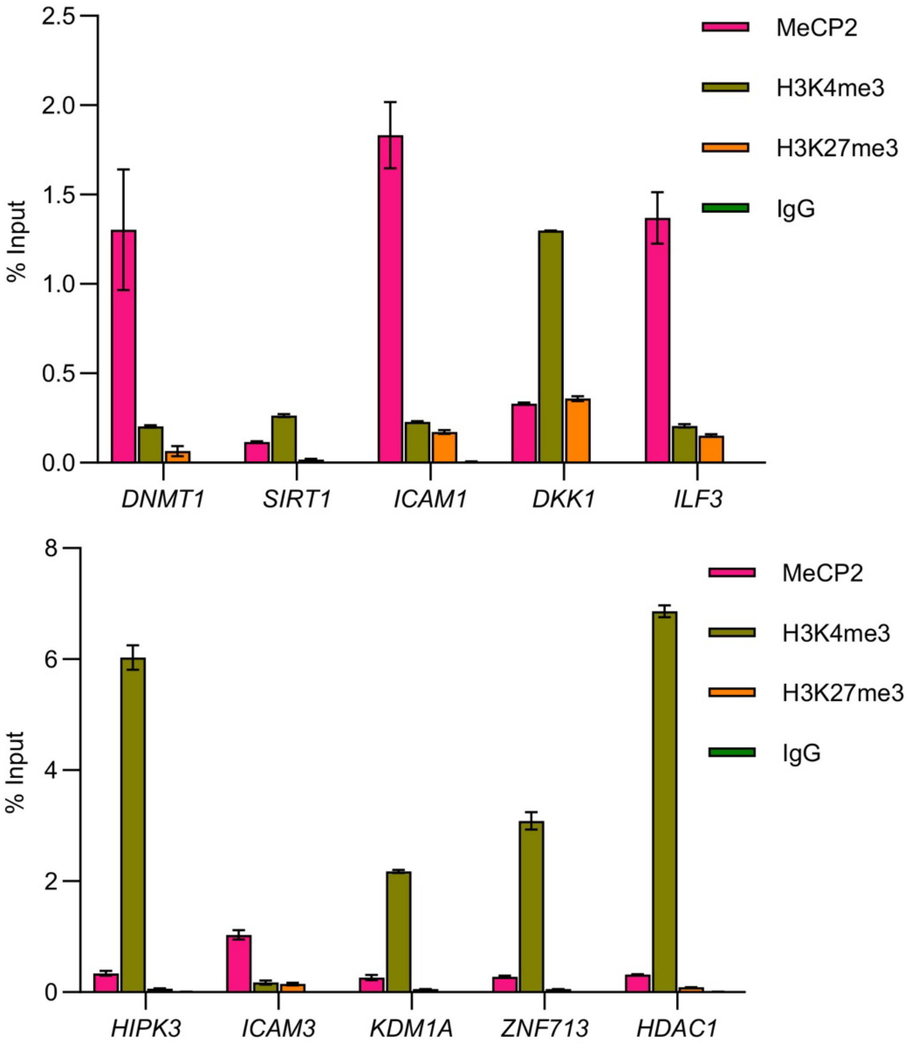


**Supplementary Figure S9.** ChIP-qPCR analysis of immunoprecipitated DNA to validate the presence of MeCP2 and histone marks, H3K4me3 and H3K27me3 in the indicated genes. Data are presented as mean ± SD (n = 3).

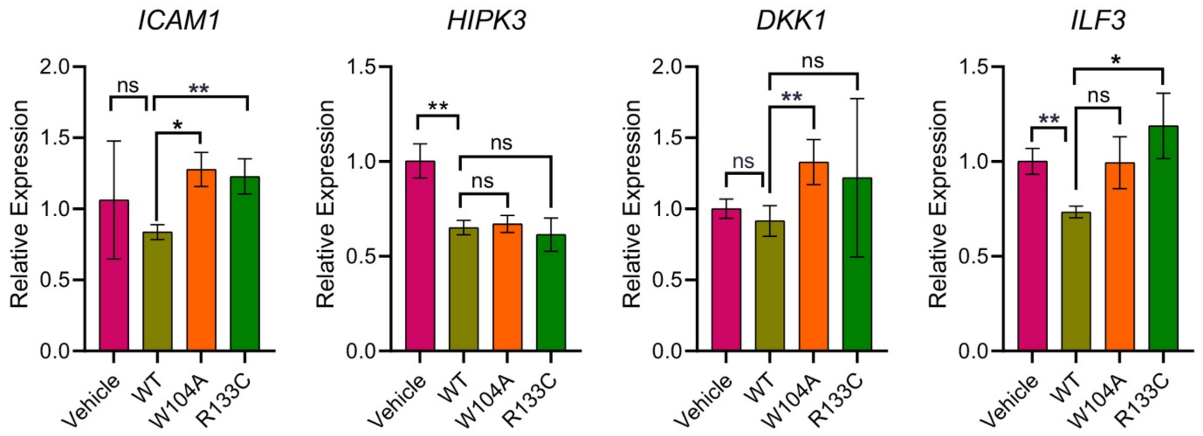


**Supplementary Figure S10**. qRT-PCR analysis of MeCP2 target genes in the presence of MeCP2-WT and its mutants W104A, and R133C. Data are presented as mean ± SD (n = 3). Statistical significance was determined using Student's t-test (*$p \leq 0.05$; **$p \leq 0.01$; ***$p \leq 0.001$; ns, not significant)

**HPLC REPORT**

| | | | |
|---|---|---|---|
| Sample: | Pep-430 APRKQLATKAARK(me3)SAPATGGVK | | Analyzed date: 30-08-2022 |
| Analyst: | Dr.RS-SBio | | |
| Column: | Gemini-NX 5µ C18 110A, 4.6*250mm | | |
| Solvent A | A: 0.1% Trifluoroacetic Acid in 100% Acetonitrile | | |
| Solvent B | B: 0.1% Trifluoroacetic Acid in 100% Water | | |
| Gradient: | | A | B |
| | 0.0min | 5% | 95% |
| | 25.0min | 30% | 70% |
| | 25.1min | 100% | 0% |
| | 30.0min | | Stop |
| Volume: | 20µl | | |
| Wavelength: | 220nm | | |
| Flow rate: | 1.0ml/min | | |

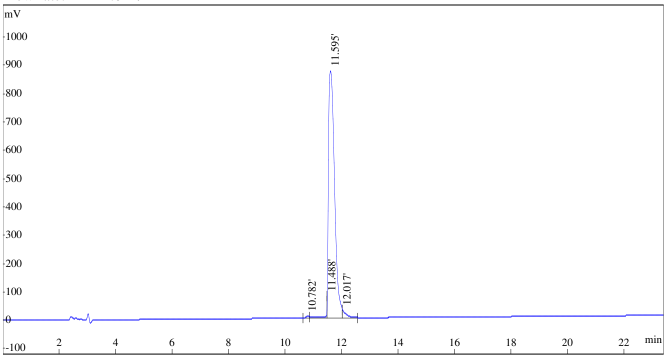


| Rank | Time | Conc. | Area | Height |
|---|---|---|---|---|
| 1 | 10.782 | 0.3655 | 46612 | 5840 |
| 2 | 11.488 | 1.1730 | 149603 | 76650 |
| 3 | 11.595 | 96.6699 | 12328976 | 868968 |
| 4 | 12.017 | 1.7916 | 228489 | 27967 |
| Total | | 100 | 12753680 | 979425 |

**Supplementary Figure S11.** HPLC purity trace for the H3K27me3 (15-36) peptide.

**MASS SPECTROMETRY REPORT**

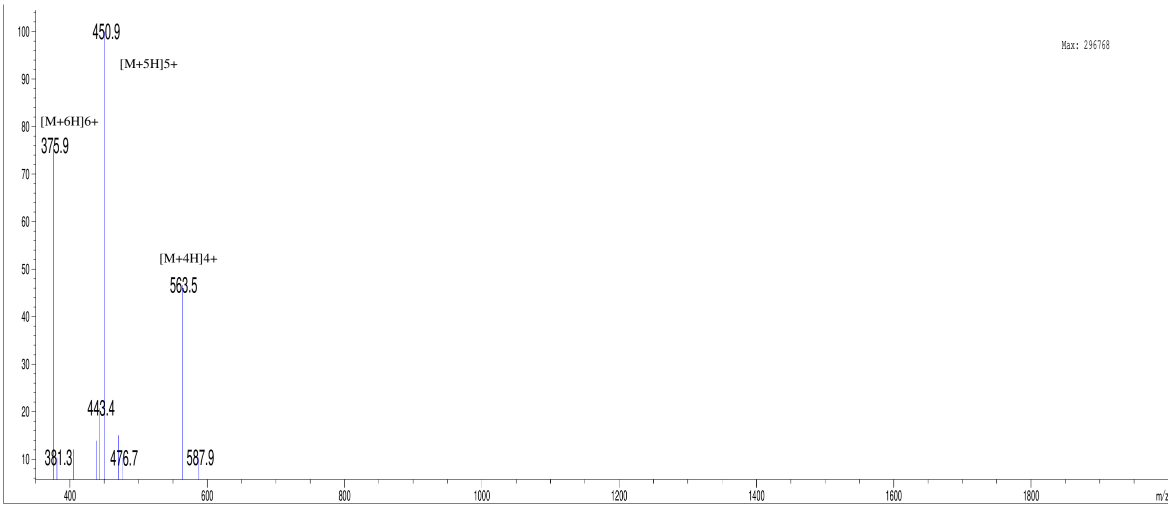


| | | | | | |
|---|---|---|---|---|---|
| Sample Description | | Instrument | Agilent-6125B | | |
| Analyzed date: | 31-08-2022 | Probe: | ESI | Probe Bias: | +4.5kv |
| Analyst: | Dr.AR-SBio | Nebulizer Gas Flow: | 1.5L/min | Detector: | 1.5kv |
| Sample: | Pep-430 APRKQLATKAARK(me3)SAPATGGVK | CDL: | -20.0v | T. Flow: | 0.2ml/min |
| M.W.: | 2249.57 | CDL Temp.: | 250 °C | B. Conc.: | 50%H2O/50%ACN |

**Supplementary Figure S12.** MS spectra for the H3K27me3 (15-36) peptide.
